# Identification and Quantification of Within-Burst Dynamics in Singly-Labeled Single-Molecule Fluorescence Lifetime Experiments

**DOI:** 10.1101/2022.05.23.493026

**Authors:** Paul David Harris, Eitan Lerner

## Abstract

Single-molecule spectroscopy has revolutionized molecular biophysics and provided means to probe how structural moieties within biomolecules spatially reorganize at different timescales. There are several single-molecule methodologies that probe local structural dynamics in the vicinity of a single dye-labeled residue, which rely on fluorescence lifetimes as readout. Nevertheless, an analytical framework to quantify dynamics in such single-molecule single-dye fluorescence bursts, at timescales of microseconds to milliseconds, has not yet been demonstrated. Here, we suggest an analytical framework for identifying and quantifying within-burst lifetime-based dynamics, such as conformational dynamics recorded in single-molecule photo-isomerization related fluorescence enhancement. After testing the capabilities of the analysis on simulations, we proceed to exhibit within-burst millisecond local structural dynamics in the unbound α-synuclein monomer. The analytical framework provided in this work paves the way for extracting a full picture of the energy landscape for the coordinate probed by fluorescence-lifetime based single-molecule measurements.

## Introduction

The advent of single-molecule fluorescence spectroscopy (SMFS) has been a boon to structural biology, allowing conformations to be directly observed, rather than the ensemble average of a large group of molecules with multiple unsynchronized conformations.^1,2^ There are a variety of SMFS modalities, which vary in time resolution^2^, and all rely on attaching fluorophores to a biomolecule, such as a protein or nucleic acid under study. These methods vary both by whether a confocal or wide-field microscope is used, and by what fluorescence phenomenon is exploited, such as Förster resonance energy transfer (FRET), photo-induced electron transfer (PET),^3,4^ or photo-isomerization related fluorescence enhancement (PIFE; commonly known as protein-induced fluorescence enhancement)^5–10^ among others.

Confocal-based methods provide the highest time resolution, using detectors sensitive to the arrival of individual photons, and typically record interphoton times on the order of a few microseconds^11,12^. In these experiments, single molecules traverse the confocal volume, and while doing so undergo multiple excitation-emission cycles. This results in periods of high photon count rate, referred to as photon bursts, which typically last a few milliseconds. In traditional burst analysis, the photon detection events of each burst are aggregated to form a single data point per burst. This is ideal for biomolecules that undergo dynamic transitions at timescales slower than milliseconds, termed *between-burst dynamics*. However, when dynamic transitions occur in timescales on the order of burst durations or faster, termed *within-burst dynamics*, molecular subpopulations blur together and require alternative techniques that analyze the photon stream directly to uncover the dynamics hidden in the blurred burst parameter histograms.

Methods such as probability distribution analysis (PDA),^13^ burst variance analysis (BVA),^14,15^ and others^2^ exist to quantify within-burst dynamics in confocal-based single-molecule FRET experiments, relying on the fundamentally centrally distributed ratiometric parameter of the FRET efficiency. In other modalities that rely on fluorescence from singly-labeled biomolecules, such as PIFE, the readout is either the fluorescence detection rate or the fluorescence lifetime, a parameter which distributes exponentially. At the present time, such single-dye single-molecule confocal-based modalities lack a method for quantifying fluorescence lifetime-based within-burst dynamics.

Out of several modalities that introduce excited-state fluorescence modulation (i.e., either quenching or enhancement), PIFE is of particular interest because (i) it only requires a single dye, simplifying labeling procedures and minimizing the potential structural perturbation that dye labeling may introduce to the biomolecule under study, and (ii) the fluorescence intensity and the fluorescence lifetime of the dye has been shown to be monotonically related to proximity of the dye to the biomolecular surface in the 0-3 nm range^5,8,16–18^, among other potential reasons that affect the dye fluorescence lifetime. This renders PIFE not only an attractive method on its own, but also a nice complement to FRET, having sensitivity to distances smaller than the scale of distance sensitivities in FRET.

Using this effect, we were able to observe different groupings of photon bursts of freely-diffusing sulfo-Cy3 (sCy3)-labeled biomolecules based on the mean value of all photon nanotimes (i.e., mean nanotime), the photon detection times relative to the moments of excitation that led to these photon emission events. Bursts with short mean nanotimes are associated with biomolecular structure having minimal steric hindrance on the sCy3-labeled residue, while those with longer mean nanotimes are associated with biomolecular structure having a relatively restricted steric environment in its vicinity. This assessment, however, still relied on assessing single-molecule bursts over an integrated parameter, and therefore the possibility of within-burst dynamics could not be precluded.

Therefore, we aspired to develop a method to quantify within-burst dynamics in single-dye single-molecule fluorescence lifetime-based measurements such as smPIFE. For this, we sought to adapt the multi-parameter photon-by-photon hidden Markov modeling (mpH^2^MM) framework^19^ that we recently introduced to function with photon nanotime data. We therefore introduce a simple and widely implementable scheme to transform the exponential distribution of photon nanotimes into set of tractable parameters, amenable for use with mpH^2^MM.

In this work we first use simulations to assess the applicability and limitations of our scheme to find within-burst dynamics using photon nanotimes. We then apply it to smPIFE data of freely-diffusing unbound α-synuclein monomer, and are able to quantify the transition rates between two conformational subpopulations.

## Results

### The divisor approach

Single photon counting data can include the delay between the moment of excitation and photon detection, a value called the photon nanotime. The maximum time measured corresponds to the laser repetition rate, creating a time window for potential photon nanotime values. Detectors typically measure the nanotime with a precision in the tens of picoseconds. When a large number of photons are available, the histogram of photon nanotimes can be used to build a fluorescence decay (Fig. 1, d). Unfortunately, a single burst typically contains far too few photons to allow for decay fitting. Therefore, alternative methods such as maximum likelihood estimation (MLE)^20^ or the phasor approach^21^ are used to determine the fluorescence lifetime of small numbers of photons. Even these methods, however, have no way to distinguish between a transition within a burst and a single state with a multiexponential decay.

**Figure 1.**
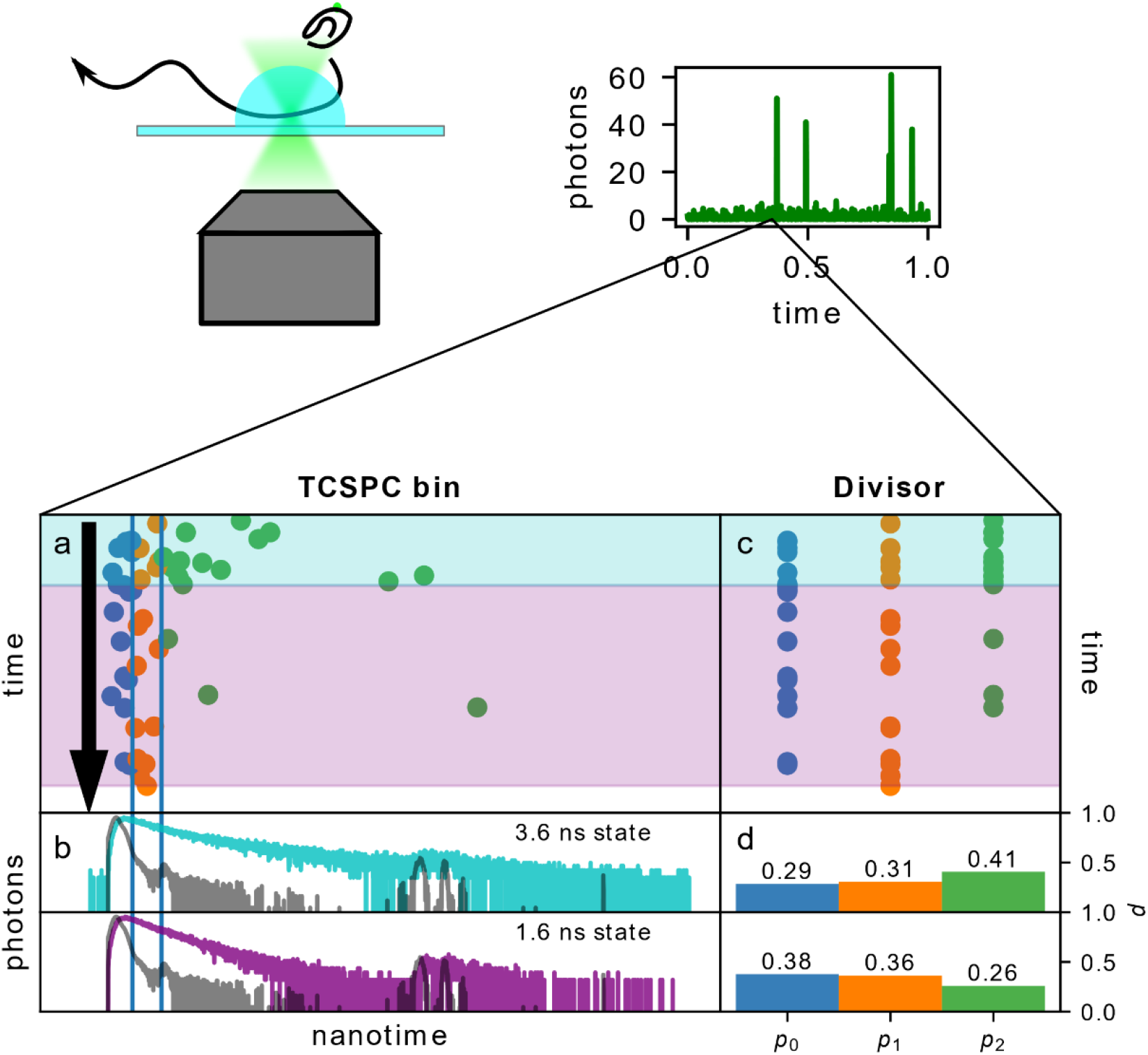
The divisor-based approach to analyzing within-burst fluorescence lifetime dynamics: a) simulated burst undergoing a transition. The absolute arrival time is represented on the vertical axis, and the photon nanotime on the horizontal. The vertical lines indicate divisors separating photons into bins. b) The simulated decays of the states in a are given in colored lines. The IRF given in grey lines. c) The same burst, but with photons organized by bins, not nanotimes. d) The probability of a photon arriving in each divisor for the two states.

MpH^2^MM is an analytical method that extends classical hidden Markov modelling to the domain of single-molecule measurements. It has previously been used to identify within-burst dynamics originating from both conformational and photophysical dynamics in smFRET experiments^19^. As such, it excels at distinguishing between data containing a mixture of states, and data with a single state of more mixed nature.

Yet, raw nanotime data is not directly applicable to mpH^2^MM, as mpH^2^MM can assess photons in single molecule data as arriving in a finite set of channels, with the relative probability of a photon arriving in a given channel to be non-negligible. Therefore, we bin the photon nanotimes based on a set of divisors (Fig 1, vertical lines in a, b, are divisors, bins shown in c, d). If two bins separated by one divisor are defined, the system mirrors the behavior described in Kim *et al*.’s work, where they describe a parameter termed *promptness ratio*^22^. Adapting equation 2 from Kim *et. al*. if a state has a single exponential decay, the lifetime can be described by Eq. 1:

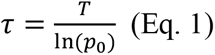

where τ is the fluorescence lifetime, *T* is the time between the laser pulse time and the divisor, and *p*_0_ is the fraction of photons detected with nanotimes smaller than the divisor. Importantly, the researchers noted in this work that the sensitivity to different fluorescent lifetimes and to multiexponential decays was dependent on the choice of the divisor, therefore the same can be expected in our analysis. Adding more divisors, we gain more sensitivity to changes between different lifetime values. However, this also reduces the probability of a photon arriving in a given bin, which reduces mpH^2^MM’s ability to identify states. Therefore, a balance must be found. Since lifetime values can vary depending on the experimental system, we attempt to develop a general approach for assigning divisors. Each method we term a *divisor scheme*. We tested a total of eight different divisor schemes, the details of which are described in the methods.

### Simulations

Using PyBroMo^23,24^, we simulate fluorescently-labeled single-dye diffusing biomolecules transitioning between two fluorescent states with two distinct fluorescence lifetimes at varying rates from as slow as 10 s^-1^ to as rapid as 10,000 s^-1^. The fluorescence quantum yield value of each state scales linearly with the fluorescence lifetime of each state. Therefore, these conditions simulate single-dye fluorescence-lifetime-based experiments with fluorescence modulation solely due to excited-state modulation (e.g., excited-state quenching or enhancement), without ground-state modulations (e.g., ground-state quenching). The short lifetime of the first state is either 1.2 or 1.6 ns, and the long lifetime of the second state is either 3.2 or 3.6 ns, with all combinations of short and long lifetime simulated. The simulated single-molecule burst data is analyzed using FRETBursts^25^ to identify single-molecule bursts. For simulations where both transition rates are slow compared to burst duration (<100 s^-1^), different mean nanotime sub-populations are clearly distinguishable for each state. For faster transition rates, more bursts of intermediate mean nanotime values are present, forming a bridge between the short and long lifetime sub-populations (Fig. 2, a). This blurring of sub-populations results from transitions between states of different fluorescence lifetimes and quantum yields, while the dye-labeled biomolecule traverses the effective excitation volume. There is a bias in burst selection for bursts of the longer lifetime state. This is due to the higher fluorescence quantum yield of the long lifetime sub-population, resulting in higher photon rates, which are more likely to have photon count rates above the background threshold. Thus, in single-dye fluorescence lifetime-based applications, where the fluorescence lifetime is linearly proportional to the fluorescence quantum yield (e.g., smPIFE, excited-state quenching), there will be an inherent bias in burst selection for longer lifetime bursts.

**Figure 2.**
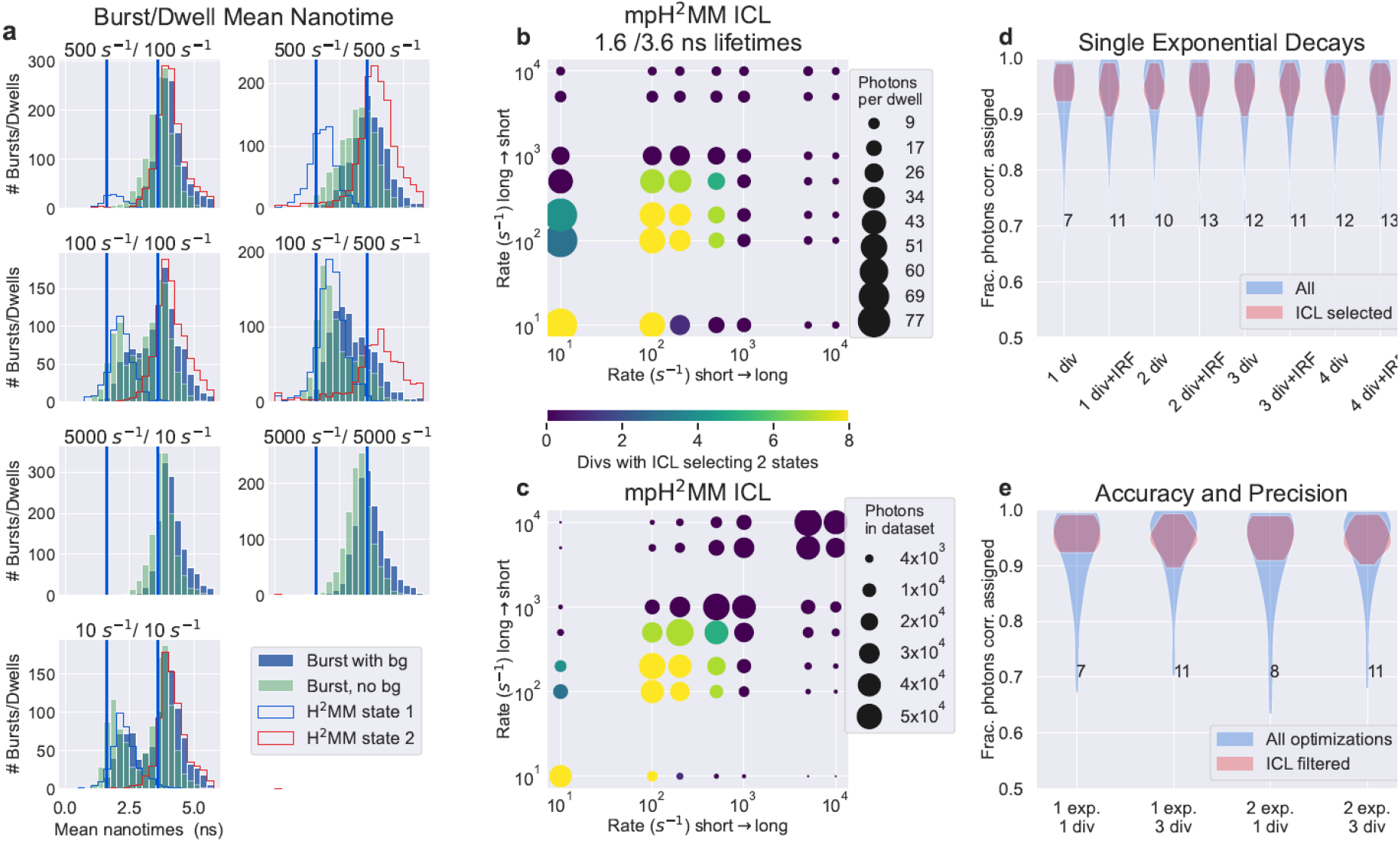
Sensitivity of mpH^2^MM in divisor-based fluorescence lifetime analysis. a) Selected examples of mean nanotime histograms of simulated data. Vertical bars indicate the ground-truth lifetime values. Filled bar histograms are mean nanotimes of bursts, blue of data including simulated background, while teal bars are the same data but with background photons excluding from calculation of mean nanotimes. Stepped line histograms show mean nanotimes of dwells ins a state within a burst determined by mpH^2^MM analysis. b, c) The minimization of the statistic, ICL, and its capability to correctly identify the number of states. Color indicates the number of divisor schemes where the ICL was minimized for the ground-truth two-state models. In b) the size of the spot indicates the mean number of photons in a dwell within a burst, for the state with the least number of photons per dwell. In c) the size of the spot indicates, for the less populated state, how many photons from that state were available across all bursts. d,e) Violin plots of the fraction of photons whose most-likely state from mpH^2^MM analysis determined by the Viterbi algorithm, matched that of the state known by the ground-truth. In d) different divisor schemes for simulations using single exponential decays are compared. In e) different divisors and simulations with mono- and bi-exponential decays are compared. Numbers indicate the number of mpH^2^MM models of the given divisor/decay combination whose ICL was minimized for the ground-truth two-state model.

Before quantitatively analyzing the data using the mpH^2^MM approach, we tested photon statistics methods that help indicate on the presence or absence of within-burst dynamics, such as burst variance analysis (BVA)^14^ and two-channel kernel-based density distribution estimator (2CDE)^26^. Our tests have shown that these approaches are not useful for analyzing within-burst dynamics in single-dye SMFS data (see detailed discussion in the SI and Figs. S1, S2).

Then, we perform mpH^2^MM analyses of all burst photon data by first choosing a divisor scheme to index the burst photon data, and then running mpH^2^MM on the indexed data. The final output per each divisor scheme can then be compared. Then, we compare the results of the mpH^2^MM analyses of the simulated data to the known ground-truth parameter values.

Analyzing data with mpH^2^MM, the first challenge is discriminating over- and under-fit models from the ideal number of states. Similar to FRET-based applications, we find that the ICL is a good statistical discriminator. We find that the ICL selects the correct number of states in most cases (Fig. 2, b, c; Figs. S3-S6). The two key limitations being that the number of consecutive photons in a given state must be sufficient (Fig. 2, b; Figs. S4-S6), and there must be a sufficient number of instances of a given state for it to be detectable (Fig. 2, c; Figs. S4-S6). The ICL was rarely minimized for three-state models, indicating that the ICL rarely selects an over-fit model (Table S1). These over-fit models all show features of over-fitting that make it possible for the user to screen for such models a simple task (Fig. S7).

To assess the influence of the size of the dataset, we also analyzed truncated versions of our simulated data. In most cases the ICL remained minimized at nearly identical results (Figs. S8-S11).

Then we assess the accuracy of the *Viterbi* algorithm, which finds the most-likely state of each photon given the optimized model and the recorded photon data. Since the simulation records the state of the molecule for each photon detection time, we could compare this state with that determined by the *Viterbi* algorithm. We found that the *Viterbi* algorithm was ∼95% accurate in assigning the correct photons to their underlying states (Fig. 2, d, e). For states represented by fluorescence decays with a single fluorescence lifetime component, all divisor schemes we tested have nearly identical accuracy. However, divisor schemes with more divisors were more likely to select the two-state model based on ICL, while the ICL of divisor schemes with fewer divisors were more likely to select under-fit single state models. Since in many cases practical systems will involve multi-component decays, it is therefore advisable to rely on divisor schemes with multiple divisors (see Table S1).

Comparing ground-truth transition rates to those detected by mpH^2^MM, we note a degree of deviation. The recovered transition rates from the short fluorescence lifetime state are consistently faster than the ground-truth values, while the recovered transition rates from the long fluorescence lifetime states are consistently slower (Fig. 3, a, b; Figs. S12-S14, repeats of this figure for other lifetime combinations, Fig. S15 for more detailed summary). This deviation is likely due to the proportionally more photons emitted from the long lifetime state per unit time compared to the short fluorescence lifetime state, biasing the results towards the long fluorescence lifetime state. In all cases the deviation was still within a factor of three at most compared ground-truth transition rates can be considered comparable, and thus mpH^2^MM extracts reasonable dynamics so long as transition rates are within an order of magnitude of the burst duration.

**Figure 3.**
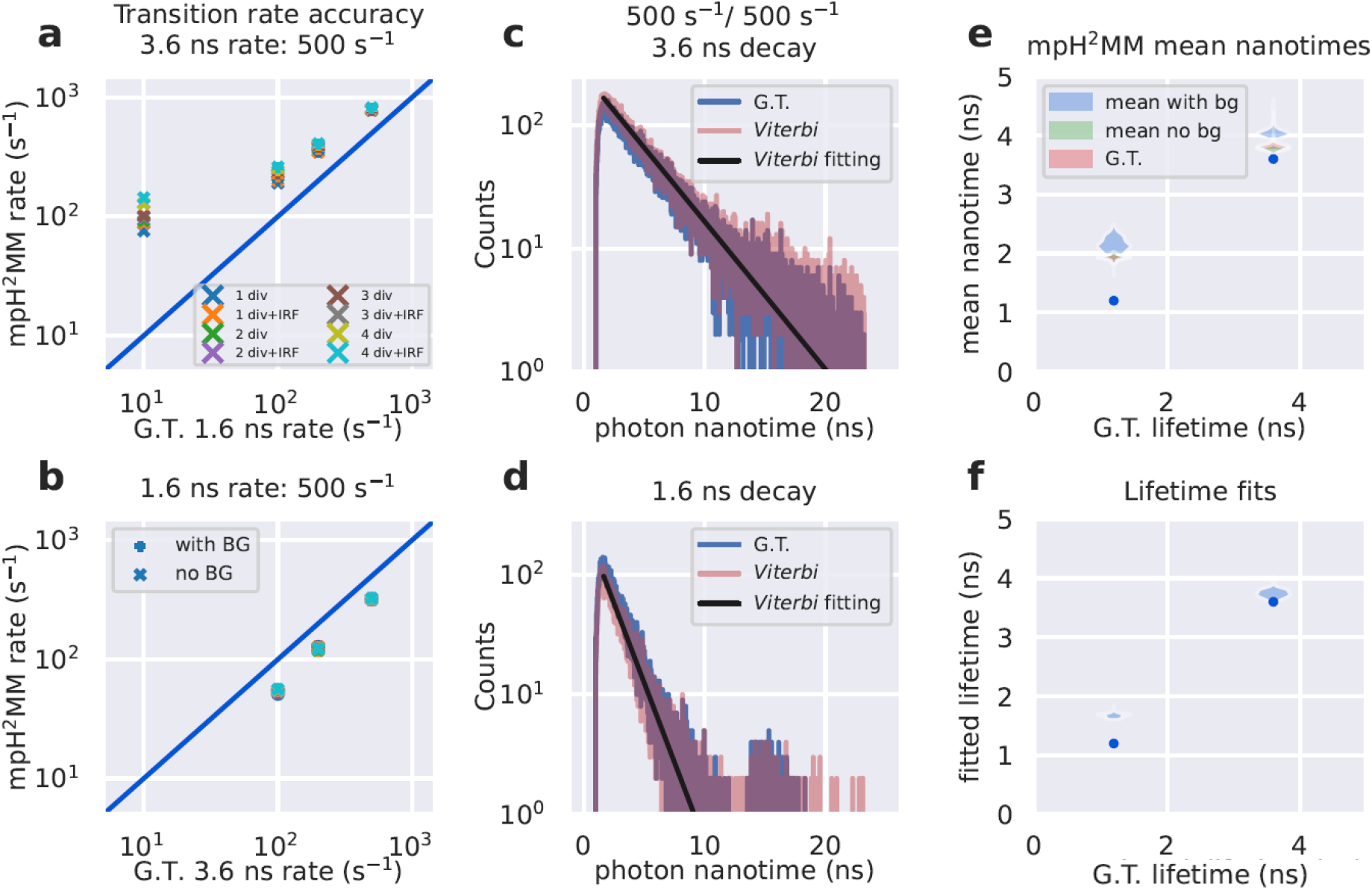
Assessment of mpH^2^MM in determining transition rates and lifetimes. a, b) the ground truth transition rate versus mpH^2^MM. Horizontal axis compares transition rate from the a) 3.6 ns or b) 1.6 ns states. In all panels, the transition rate from the other state is held constant at 500 s^-1^. + signs indicate mpH^2^MM optimizations with all photons, while x signs indicate optimizations where background photons were excluded. c, d) comparison of fluorescence decays of the ground-truth (blue) and reconstructed based on most-likely state derived from the *Viterbi* algorithm employed on the results of mpH^2^MM analyses of the simulated data (red). Black line shows biexponential fitting of *Viterbi* based decay. Both are for simulations with 500 s^-1^ transition rate for both states (i.e., the transition rates are symmetric), c) shows 3.6 ns state, d) shows 1.6 ns state. e) Violin plot showing the mean nanotimes of all states using mpH^2^MM for all ICL selected models of simulations with states of 1.6 and 3.6 ns. Blue plots for data with background photons included, green for data with background photons excluded, and red of mean nanotimes derived from the ground truth. f) Violin plots of monoexponential fittings of Viterbi decays of same data s in panel e). G.T. stands for ground truth.

Finally, we turn to assessing the accuracy of fluorescence lifetimes in states assessed by mpH^2^MM. This can be assessed in multiple ways. If a single divisor is used, it can be treated using Eq. 1. However, for multi-divisor schemes multiple potential lifetimes can be calculated, and thus deriving the fluorescence lifetime is non-trivial. We therefore prefer to use the results of the *Viterbi* algorithm to assign photons to states, build fluorescent decays, and extract the fluorescent lifetime values from these decays. Further confirming the accuracy of *Viterbi*/mpH^2^MM, these decays are consistent with the ground-truth lifetimes. The extracted decays closely mirror the ground-truth (Fig. 3, b, c; Figs. S12-S14). Fitting these fluorescence decays to exponential decays, we retrieve fluorescence lifetimes that are within 0.2 ns of the ground-truth (Fig. 3, e, f). When we use multi-exponential decays specific lifetime components were less precise with fittings, however, the correspondence of the decays with the available ground-truth photons indicates that our inaccuracies were more a result of the lack of photons in each lifetime component, due to the short time of our simulations, and given larger datasets, similarly accurate lifetimes could likely be extracted.

Individual dwells in a state lack sufficient photons to construct fluorescent decays, but the mean nanotime of photons in dwells can be used. Whether dealing with bursts or dwells within bursts, the mean of the distribution is slightly larger than the simulated ground-truth lifetime values. We found that this is the result of two primary factors: (i) background photons, which are equally probable at any point in the decay, and (ii) the exclusion of the IRF. When background photons are excluded, mean nanotimes decrease, but are still longer than the ground-truth values of the simulation (Fig. 3, e).

Thus, our simulations demonstrate that mpH^2^MM is capable of disentangling within-burst dynamics in single-dye fluorescence lifetime data. While some systematic bias favors states with higher intrinsic brightness, the fundamental recovery of both lifetime states and the transition rate data is sound.

### SmPIFE-based millisecond transitions in unbound α-synuclein monomer

After testing the applicability of the multi-divisor approach and mpH^2^MM on simulations, we demonstrate the usefulness of this approach on an important biomolecular system. α-Synuclein (α-syn) is an intrinsically disordered protein that self-associates into oligomers, aggregates and amyloid fibrils, implicated in the molecular etiology of Parkinson’s disease, among other neurodegenerative diseases^27–29^. Upon binding to membranes or self-associating α-syn can gain partial folded structures^30–37^. However, when unbound, α-syn is mostly unstructured^38^ and exhibits rapid (within hundreds of nanoseconds) conformational dynamics, as has been shown in the analyses of single-molecule FRET measurements^32,39^. These important findings were focused on the dynamics of the distance between pairs of dye-labeled residues far apart on the main chain, using the 3-10 nm distance sensitivity of single-molecule FRET^2^. These results clearly report on the disordered nature of nonlocal interactions in the unbound α-syn. To better understand whether this also applies to local interactions, we decided to perform smPIFE measurements of different dye-labeled residues within the unbound α-syn, where according to the previous findings from FRET, we did not expect to observe more than a single time-averaged population of fluorescence lifetime values.

Indeed, we have recently used smPIFE, among other methods, to investigate the conformational dynamics of the unbound α-synuclein monomer.^40^ Histograms of burst-based mean nanotimes indicated that there are at least two interconverting α-syn sub-populations, each with distinct mean values. Following that, burst recurrence analysis of single particles (RASP)^41^ indicated a signature of transition dynamics between these sub-populations in the millisecond time-scale. RASP, however, reports on between-burst dynamics, which represents timescales longer than the ones that may occur within bursts. We therefore applied our divisor approach with mpH^2^MM to this data to see if within-burst dynamics could be detected, and perhaps the transition rates quantified.

We test this on the unbound α-syn monomer singly-labeled with sCy3 at residues 26, 56 and 140. Across multiple divisor schemes, the ICL predicts two states. While two states are consistently found, the transition rates across different positions vary. At position 26, they are less than 100 s^-1^, with almost no transitions found. At positions 56 and 140, however, the transition rates were larger than that value (up to 330 s^-1^), indicating a degree of within-burst dynamics. Using the *Viterbi* algorithm to segment the data into dwells in states, we generate histograms of the mean lifetime of dwells, enabling the recovery of state-relevant sub-populations without relying on fitting the burst-based mean nanotime histograms to sums of Gaussians or other fitting functions (Fig. 4). We are then able to determine the mean nanotime of each state. For all labeling residues, these are centered at ∼1.0 and ∼2.2 ns.

**Figure 4.**
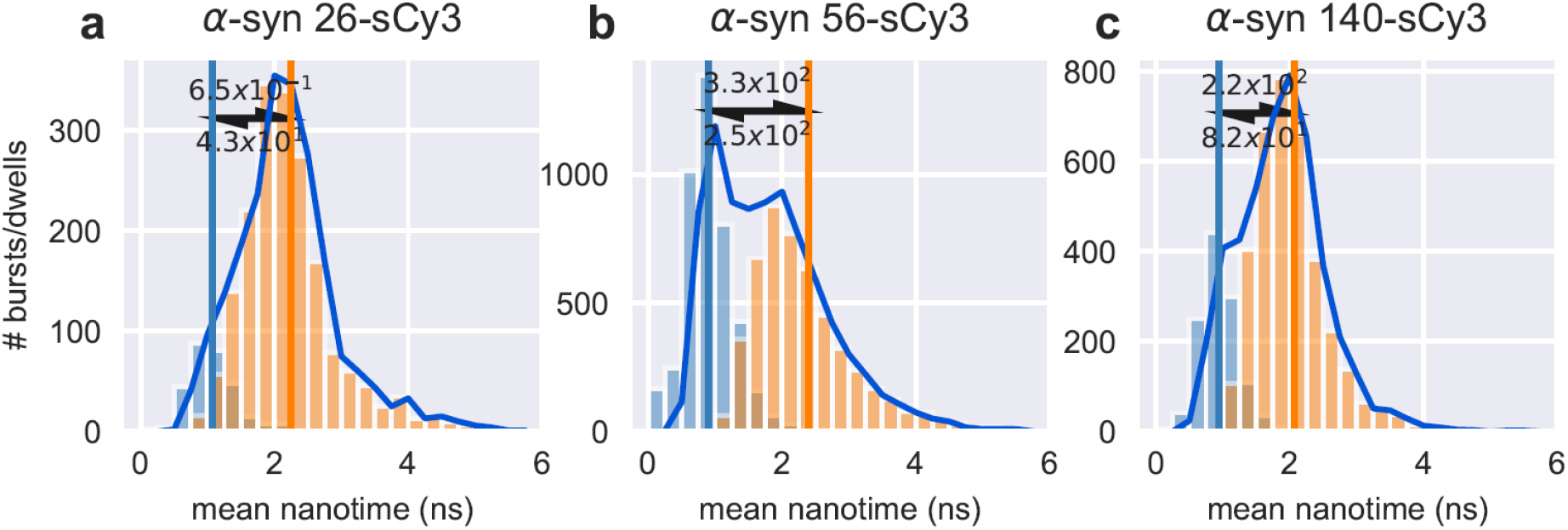
MpH^2^MM analysis of the unbound α-syn monomer labeled at positions a) 26, b) 56 and c) 140 with sCy3. Barred histograms indicate mean nanotimes of state dwells within bursts, while lined histograms indicate burst-based mean nanotimes.

Thus, we demonstrate that mpH^2^MM is capable of quantitatively characterizing transition dynamics in experimental data, such as smPIFE of the unbound α-syn monomer.

## Discussion

Using the divisor-based approach in mpH^2^MM, we are able to reduce intractably large and exponentially-distributed parameter of fluorescence nanotimes into a tractable set of parameters, enabling the detection of within-burst lifetime state dynamics using mpH^2^MM. This is especially powerful as both the fluorescence decays of states and transition rates are simultaneously recovered. This was applied in the case of smPIFE data, where the primary parameter is the mean fluorescence nanotime per state. Given a sufficiently bright sub-population, we are able to resolve transition rates and mean lifetime values reliably.

Using mpH^2^MM analysis, we recover the millisecond dynamics occurring between two major states in the unbound α-synuclein (α-syn) monomer. In these results, each state has a different mean fluorescence nanotime value that can, among other factors, represent different degrees of steric restriction imposed on the sCy3 dye conjugated to a specific residue in α-syn. This, in turn, reports on millisecond dynamics in structures that locally influence the vicinity of the dye-conjugated residue. We show this millisecond local structural dynamics occurs within bursts in the vicinity of residues 56 in the edge region of the N-terminal segment of α-syn, and 140 at the terminus of the acidic C-terminal segment of α-syn. We could not recover within-burst lifetime-based dynamics in the vicinity of residue 26 in the N-terminal segment. However, the burst-based mean nanotime histograms report on multiple sub-populations, which points towards lifetime-based dynamics in the vicinity of this residue slower than burst durations. Combined with past results from smFRET-based measurements^32,39^, we can propose that α-syn is a protein that exhibits intrinsic disorder characteristics between residues far apart along the primary sequence, perhaps representing no stable nonlocal interactions, with millisecond stable local interactions that may promote stable local structural elements. Now, with the ability to identify and quantify within-burst dynamics in smPIFE measurements, and by doing so enhancing the description of dynamics in biomolecules, we believe smPIFE could be used to study many other biomolecular systems.

While the divisor-based mpH^2^MM approach has been shown here to uniquely aid in analyzing fluorescence lifetime-based within-burst dynamics, it is important to mention that the 2D-fluorescence lifetime correlation (2D-FLC) methodology^42,43^ can help in such analyses. In 2D-FLC, 2D-nanotime correlation functions are being inverse-Laplace transformed into 2D lifetime population maps. While the 2D-FLC is reported to be sensitive to within-burst dynamics as rapid as the interphoton times, hence typically microseconds, the transformation of the data might require much more photon data relative to the divisor-based mpH^2^MM approach. Therefore, we believe that while both approaches might be complementary in analyzing single-dye fluorescence lifetime-based dynamics, the divisor-based mpH^2^MM approach might fit the low number of photon nanotimes in each single molecule burst. Importantly, dynamics in the milliseconds and sub-milliseconds many times do not provide a full picture of the fluorescence fluorescence-based dynamics. In many ways, different modalities of fluorescence correlation spectroscopy (FCS) employed on freely-diffusing single molecules through a confocal spot can also report on fluorescence-based dynamics in the microseconds and even faster (see full coverage in a recent review^2^). However, these approaches might not solely report on transitions between different fluorescence-based states, but also between bright fluorescent states and dark photo-blinked states, and it might be difficult to distinguish between these different processes based on FCS of single dye fluorescence data.

In this work we tested the application of the divisor-based mpH^2^MM approach for analyzing fluorescence lifetime-based single-molecule burst data assuming that the fluorescence lifetimes are linearly proportional to the molecular brightness, and also assuming fluorescence modulation is due solely to excited-state modulation (e.g., quenching or enhancement). However, ground-state modulation, such as ground-state quenching could also exist in the data, but not represented by dynamics between fluorescence lifetime states. At the extreme case, such ground-state modulation that leads to fluorescence dynamics will be seen as dynamics between two states with different brightness values and no difference in fluorescence lifetime values. Therefore, to shed light on these additional possibilities, we suggest data treatments as were previously suggested by Kondo *et al*.^43^, namely to present also the brightness of each dwell or burst where a single molecule was identified as being in a given lifetime state, in a 2D brightness versus lifetime map. In such a map, subpopulations off the diagonal represent a fraction of ground-state and excited-state modulation dynamics.

While the divisor-based approach to integrating photon nanotimes into mpH^2^MM is typically applicable to PIFE, it is also possible to apply it to other modalities that include photon nanotimes, such as with single-molecule FRET. However, since FRET efficiencies and fluorescence lifetimes are intrinsically linked, barring additional complicated protein interactions, this also has the potential to introduce redundant data. Nevertheless, there are situations in which the ratiometric FRET efficiency parameter values differ from the FRET efficiency parameter values calculated from donor fluorescence lifetimes. One such case is when the doubly-labeled system exhibits within-burst donor-acceptor distance dynamics that can even be faster than the typical inter-photon times. This deviation occurs due to the dynamics, where the high FRET state has not only lower donor lifetime than the lower FRET state, but also less donor fluorescence photons - a situation analogous to the changes in both fluorescence lifetime and quantum yield in PIFE. Understanding the sources of these differences paved the way for Seidel and co-workers to develop an analytical framework dubbed FRET-Lines^44^ to retrieve the underlying FRET dynamics from multi-parameter fluorescence detection (MFD) time-resolved smFRET measurements^45–47^. In parallel to this approach, we provide the mpH^2^MM approach for analyzing and quantifying within-burst dynamics in such confocal-based single-molecule measurements that refer to ratiometric FRET efficiency, donor and acceptor fluorescence lifetimes, fluorescence anisotropies and other parameters. However, note that we currently do not gain any additional useful information from applying the MFD approach compared with our previously implemented version of mpH^2^MM, if not the donor nor the acceptor are undergoing changes independent of FRET. Nevertheless, there are modalities, such as the combined PIFE-FRET^17^, in which this analysis can be useful, to decouple three types of within-burst dynamics: (i) FRET dynamics, (ii) PIFE dynamics, and (iii) photophysical dynamics. In fact, retrieving the mean values from mpH^2^MM analyses of PIFE-FRET could help in linking these values to their underlying physical meanings by using the existing theoretical framework of PIFE-FRET^8^.

Incorporating more parameters combats the limited information content of single-molecule methods, which favor high time resolution, at the expense of spatial information. With more parameters available in analysis, the greater is the ability to resolve and predict the unique properties of each conformational subpopulation. The divisor approach removes one more limitation by allowing any parameter, including ones that do not centrally distribute, to be converted into a tractable ratiometric parameter that can then be used in data analyses such as mpH^2^MM. This, in turn, brings us one step closer to a full picture of the microstates of biomolecules.

## Experimental Methods

### SmPIFE measurements of sCy3-labeled α-Syn variants

The whole preparative procedure, from the expression, through the dye labeling and the purification were explained in full by Zaer and Lerner^18^.

All smPIFE measurements and burst analyses of α-syn were performed and analyzed exactly as recently explained in Chen and Zaer *et al*.^40^.

### PyBroMo Simulations

In PyBroMo, 20 particles in a 8×8×12 μm^3^ box are simulated with a diffusion constant of 12 μm^2^/s. The time step of the simulation was set to 500 ns. Once single-molecule diffusion trajectories were built, then PyBroMo simulated the macrotime of photon detections. The point spread function (PSF) numerically solved and provided by PSFlab^48^ is used to calculate the photon detection rate of each simulated single molecule at each moment, with different photon rates at the center of the PSF, better known as the molecular brightness values per each state. Then, the instantaneous photon rates are used for sampling detected photons out of the Poisson distribution with these photon rates. Then, background photons are added using a background count rate of 500 counts/second. Four different molecular brightness values were simulated for each molecule, one for each of the possible states of which two will be selected once the final trajectory is built. Since molecular brightness and lifetime in fluorescence lifetime-based applications such as PIFE have been shown to linearly depend on each other, a universal base molecular brightness and base fluorescence lifetime are set as 275,000 counts/second, and 4.0 ns respectively. Then, ratios of 0.3, 0.4, 0.8, and 0.9 are set as degrees of effects on reducing the molecular brightness and the fluorescence lifetime. Multiplying the base value by the ratio leads to the actual value used in each simulation. For the simulation of states with multi-exponential fluorescence decays, an additional decay with a base lifetime of 1.2 ns is incorporated. This shorter lifetime component is also scaled with the ratio of the brightness and lifetime reduction effect. This shorter lifetime component contributes 10% of the photons in the simulations of states with multi-exponential fluorescence decays.

PyBroMo simulates photon detection times. However, it does not yet simulate either photon nanotimes or dynamic transitions between lifetime-based states. Therefore, these effects were added through additional layers after PyBroMo simulations. To simulate photon nanotimes, a Monte-Carlo simulation is used, simulating an exponential fluorescence decay, and then adding an additional delay pulled from the experimental IRF of our setup, described in Zaer and Lerner^18^. Drawing on previous work, transitions between states are made by stitching photon trajectories together. First, dwell times in each state are simulated by taking random exponential distributions based on the transition rate. This forms time windows for each molecule when it is in a given state. Then, for each simulated state, photons within those time windows are selected, and stitched together to create the final photon trajectories of simulations.

### Burst Search and Selection

Both simulated and experimental data are imported into FRETBursts^25^, and background is first assessed for each time interval of 30 s. Then burst search is performed by selecting bursts with an instantaneous count rate of F=6 time the background rate, with the instantaneous count rate calculated using a sliding window of m=10 photons. Bursts are further refined to require a minimum width of 1 ms, a minimum peak count rate of 20,000 counts/second, and a minimum size of 25 photons.

This provided the base burst selection on which all downstream processing was conducted. Whenever mean nanotimes are discussed, a threshold is set denoting the end of the instrument response function (IRF). The mean delay of photons arriving after this threshold relative to said threshold is then calculated, and this is the mean nanotime. Photons arriving before this threshold are excluded from the mean nanotime analysis. This method is applied equally to burst- and dwell-based mean nanotimes and used in BVA calculations.

### MpH^2^MM analysis

In mpH2MM, photons must be assigned indices, which are assigned based on the bin into which the nanotime of the falls. We test a total of eight different divisor schemes, which assign photons to bins differently. These divisor schemes differ in how many divisors are used, and whether or not the IRF was assigned to its own bin. Thus, divisor schemes can be assigned into two groups: the non-IRF and with IRF divisor schemes. For non-IRF scheme, the divisors were set to equally divide the total photon nanotimes of all photons within bursts into bins. Thus, if there was one divisor, it would be placed at the 50^th^ percentile, if two divisors, at the 33^rd^ and 66^th^ percentiles. The IRF schemes followed a similar pattern, except that an additional divisor was set at the end of the IRF, and the percentiles were assigned according to the nanotimes of only photons which arrived after the IRF divisor. We test from one to for divisors in the non-IRF scheme, and 1 to 4 divisors (which have n+1 divisors due to the additional divisor of the IRF) in the IRF scheme. All analyses were performed using the H2MM_C python package^19^. Model optimizations were performed on single-, two- and three-state models. The *Viterbi* algorithm was then applied to determine the most-likely state path-based optimized model, and to determine the ICL of each optimization. The state path was then segmented into state dwells, on which duration, and mean photon nanotimes could be assessed. Further, using the most-likely state assignment, the photon nanotimes are assembled to create state-based fluorescent decays. Finally, for analysis of the accuracy of the state assignment, the state assigned by *Viterbi* is compared to the ground-truth of the simulation. For this comparison, photons are assigned as correct, incorrect, or not applicable, when the ground-truth origin of the photon was either background, or from a molecule that did not contribute the majority of photons in the burst. In this final case, the result of applying the *Viterbi* algorithm is ignored. For the assessment of the fraction of photon correctly assigned to a state, these background photons are excluded. For mpH^2^MM analysis of α-syn data, optimizations are carried out for increasing numbers of states until the ICL ceases to decrease.

## Acknowledgements

We would like to thank Drs. Asaf Grupi, Dan Amir, and Elisha Haas from the Mina & Everard Goodman Faculty of Life Sciences in Bar Ilan University for sharing the plasmids of αSyn bearing single cysteine mutations. This project was supported by the Israel Science Foundation (grant 1768/15 to N.K.; grant 3565/20 to E.L., within the KillCorona – Curbing Coronavirus Research Program), the NIH (grant R01 GM130942 to E.L. as a subaward), by the Milner Fund (to E.L.), and by the Hebrew University of Jerusalem (start-up funds to E.L.).

