## Supplementary Information for "Identification and Quantification of Within-Burst Dynamics in Singly-Labeled Single-Molecule Fluorescence Lifetime Experiments"

**BVA and 2CDE analysis**

Burst variance analysis (BVA)[1] and two-channel kernel-based density distribution estimator (2CDE)[2] are photon statistics-based qualitative analyses used in smFRET to test for FRET-based within-burst dynamics. These are often used to determine if further analysis with mpH$^2$MM or other similar methods is warranted. We test if these analyses can also be used in single-dye lifetime-based single-molecule fluorescence detection, prior to employing fully quantitative approaches such as mpH$^2$MM. In theory, a method analogous to BVA could work for mean fluorescence nanotimes. Unfortunately, we find that even for static populations, the short mean nanotime sub-population alone exhibits a signature that would be considered within-burst dynamics in BVA (Fig. S1). This effect can be traced to the presence of background photons, as when we remove these photons from the data set, this signature disappears. However, as background photons are inherent to single-molecule fluorescence detection, a direct removal of background photons is not feasible. Note that the background photon rate we use in these simulations is based on the background rate for hybrid photo-multiplier tubes, which are lower than the typical background rates typically attained with single-photon avalanche diode (SPAD) detectors. Therefore, BVA is not suitable for detecting within-burst dynamics in single-molecule single-dye fluorescence lifetime-based experiments. We also attempt to adapt 2CDE using the divisor approach proposed for mpH$^2$MM, but the opposite problem occurs compared to BVA: all datasets show no signatures of within-burst dynamics in 2CDE (Fig. S2).

**BVA and 2CDE implementation**

Bust variance analysis is calculated analogously to BVA in FRET[1], replacing the ratiometric FRET efficiency parameter, $E$, with the mean photon nanotime in a burst. Photons in each burst are segmented into groups of $m$ consecutive photons, and the standard deviation is calculated as in Eq. S1:

$$\sigma_\tau = \sqrt{\frac{\sum_{i=1}^{n}(\tau_i - \tau_{burst})^2}{m}} \text{ (Eq. S1)}$$

where $\tau_i$ is the mean nanotime of each group of consecutive photons, $\tau_{burst}$ is the mean nanotime of the entire burst, and $n$ is the number of groups in the burst.

Since photon nanotimes follow an exponential distribution, if the burst contains a single lifetime, the standard deviation is expected to be proportional to Eq. S2:

$$\sigma(\tau, n) = \tau/m \text{ (Eq. S2)}$$

This, obviously, influences the shape of the BVA static line to be different than its shape in BVA employed on ratiometric FRET. For plotting, we therefore choose to plot $\sigma_\tau/\tau_{burst}$ so that bursts with no within-burst dynamics will have a constant value of $\sigma_\tau/\tau_{burst} = 1/m$ .

To adapt the equations of 2CDE to fluorescence lifetime data, we took the mean nanotime as a $t_{gate}$, and treat photons with nanotimes less than $t_{gate}$ as analogous to donor photons, and photons with nanotimes greater than $t_{gate}$ as analogous to acceptor photons, in analogy to the original FRET implementation of the 2CDE approach.

**Over-fitting**

To assess the influence of the size of the dataset, we also analyzed truncated versions of our simulated data. In most cases the ICL remained minimized at the same number of states. However, for faster transition rates, it was more likely for the ICL to be minimized at single-state instead of two-states. Importantly, real data sets are usually at least twice as large as our simulated data. Therefore, the reliability of state selection and parameter values will likely exceed that which we observe in our simulated data.

### Supporting Figures and Tables

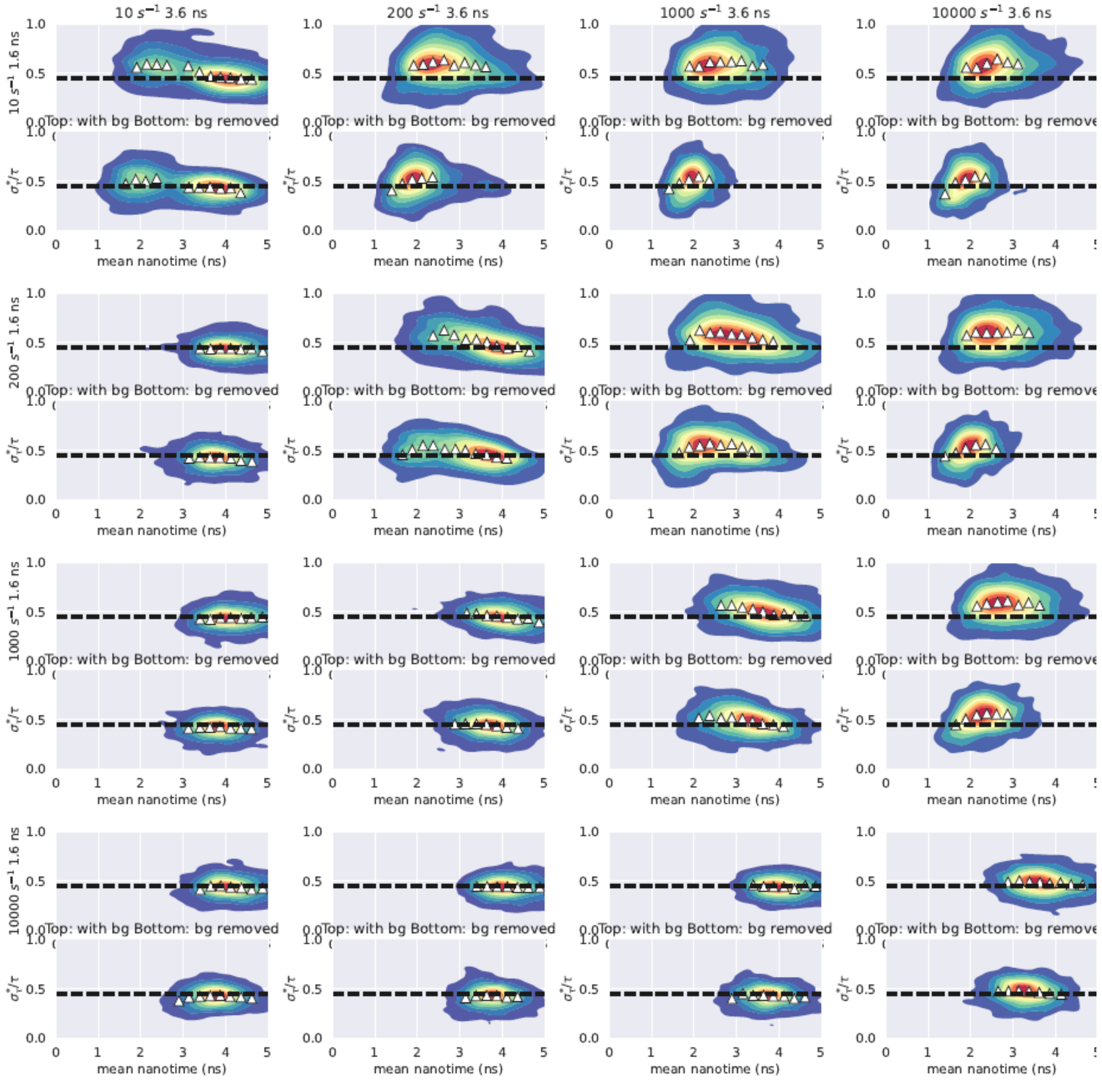

Figure S1. Examples of BVA analyses of simulated data at different transition rates. Upper panels show simulated data including background photons in BVA analysis, while lower panels are identical simulations, but with all background photons excluded from BVA analysis,

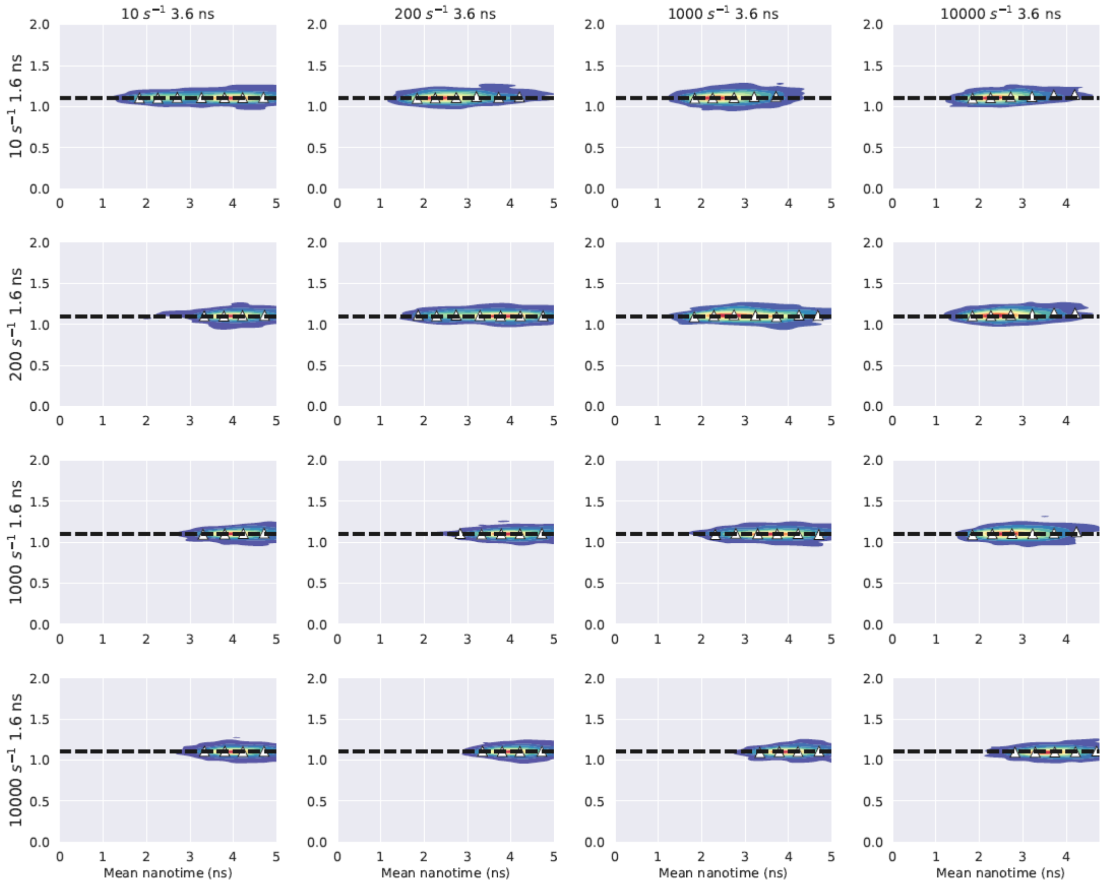

Figure S2. Examples of 2CDE analyses of bursts on simulated data. Each panel is a different simulation, transition rate from the 1.6 ns lifetime state (single exponential) increases in successive rows, while transition rate from the 3.6 ns (single exponential) transition rate increases across columns. Dotted line indicates threshold for dynamics.

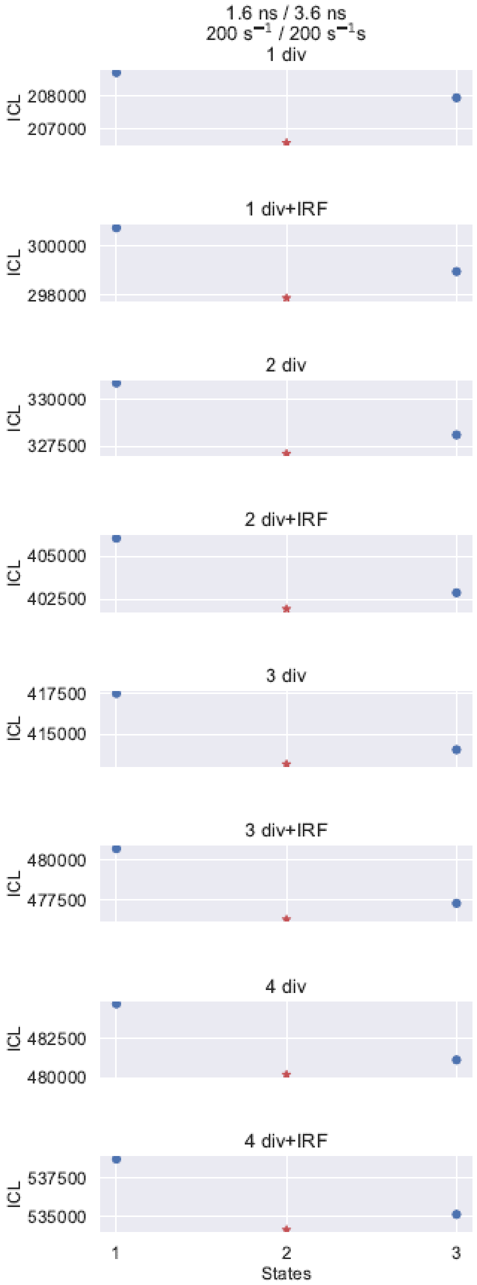

Figure S3. ICL of optimization with different divisor schemes of the ground-truth two state model with 1.6 and 3.6 ns lifetime states and 200 $s^{-1}$ transition rates. Blue circles are state models with non-minimal ICL values, while red stars indicate the state model with minimal (selected) ICL, which recovered the ground-truth state model.

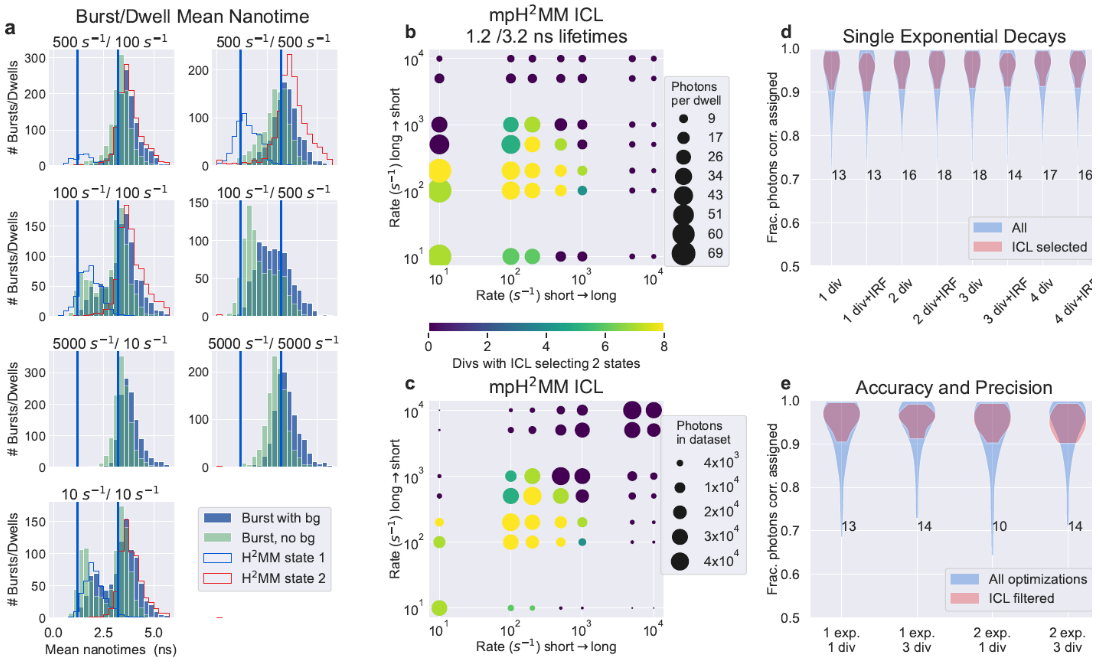

Figure S4. Model selection and accuracy for simulations with states with 1.2 and 3.2 ns fluorescence lifetimes. Panel arrangement is the same as in Fig. 2 of the main text.

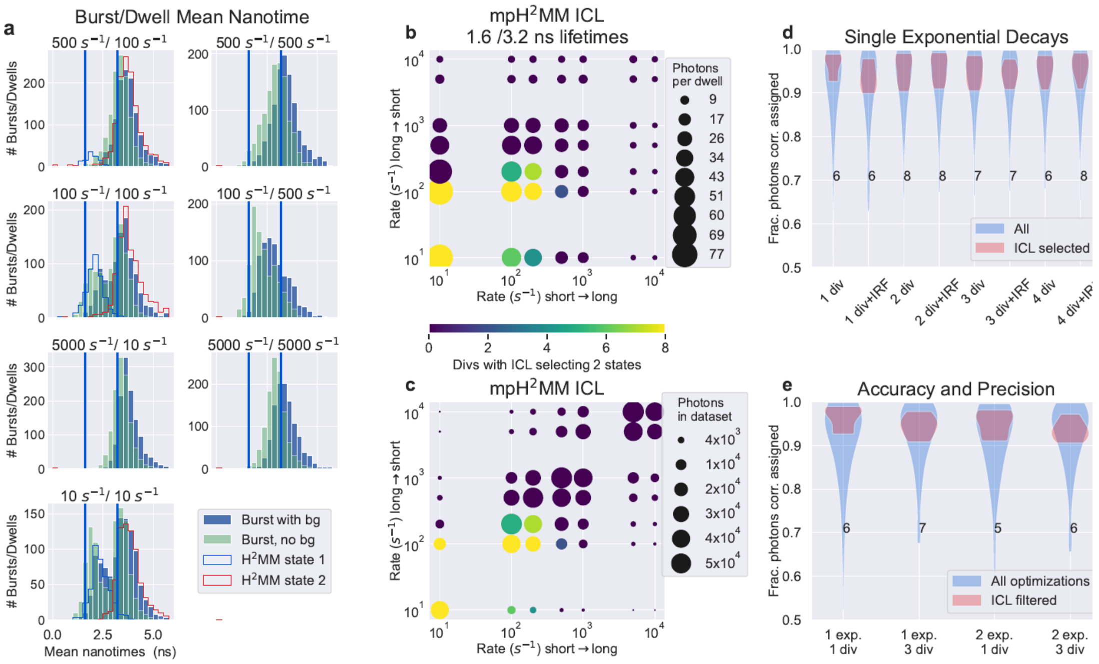

Figure S5. Model selection and accuracy for simulations with states with 1.6 and 3.2 ns fluorescence lifetimes. Panel arrangement is the same is in Fig. 2 of the main text.

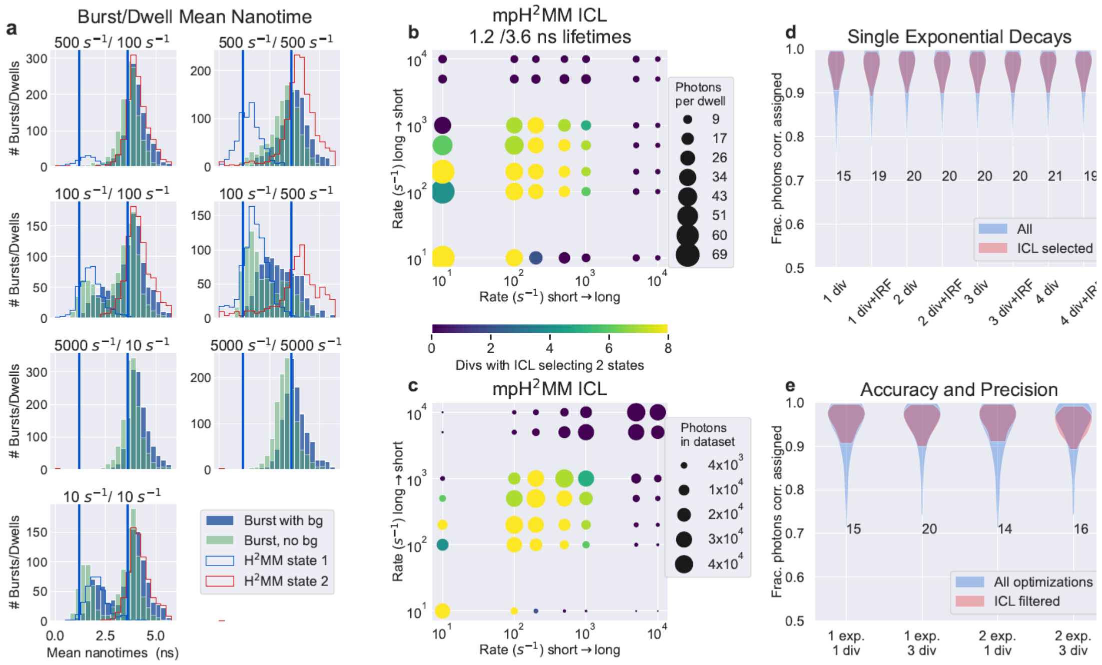

Figure S6. Model selection and accuracy for simulations with states with 1.2 and 3.6 ns fluorescence lifetimes. Panel arrangement is the same is in Fig. 2 of the main text.

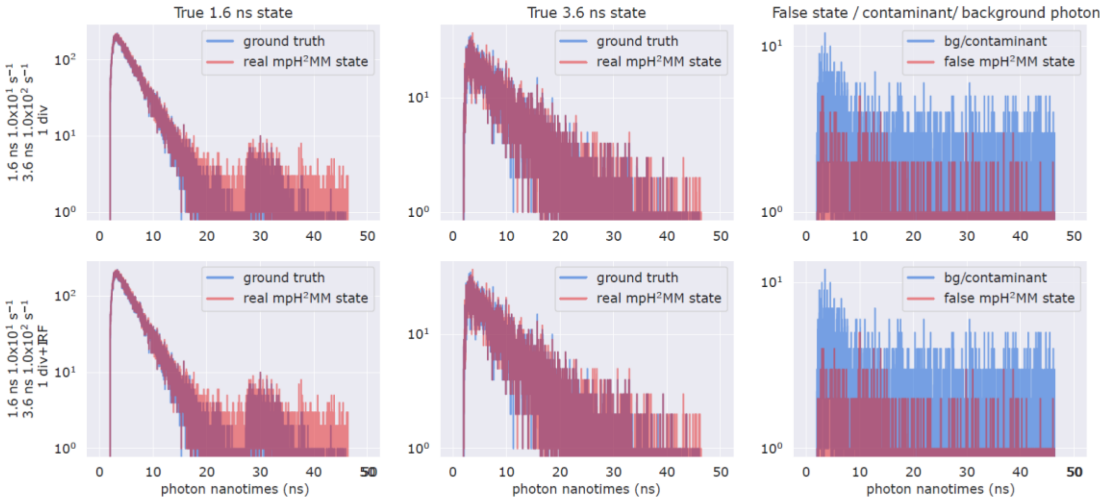

Figure S7. Two examples of mpH$^2$MM optimizations where the ICL was minimized at three states. Each row is a single data/divisor scheme combination. Each column a given state. Red lines indicate recovered decays, while blue represents the ground truth. For the third column, the blue line indicates the decays from both contaminant and background photons. Note how the third state in mpH$^2$MM models has a very small number of photons thus raising suspicion as to the reliability of the state.

**Truncated datasets**:

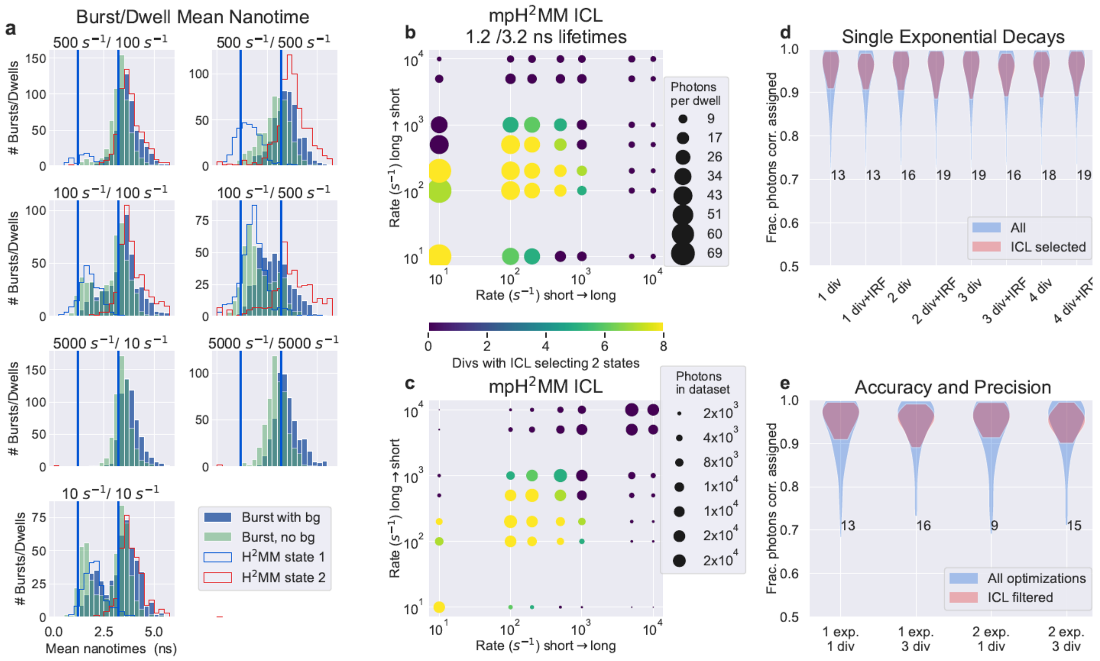

Figure S8. Model selection and accuracy for truncated simulations with states with 1.2 and 3.2 ns lifetimes. Panel arrangement is the same is in Figure 2 of the main text

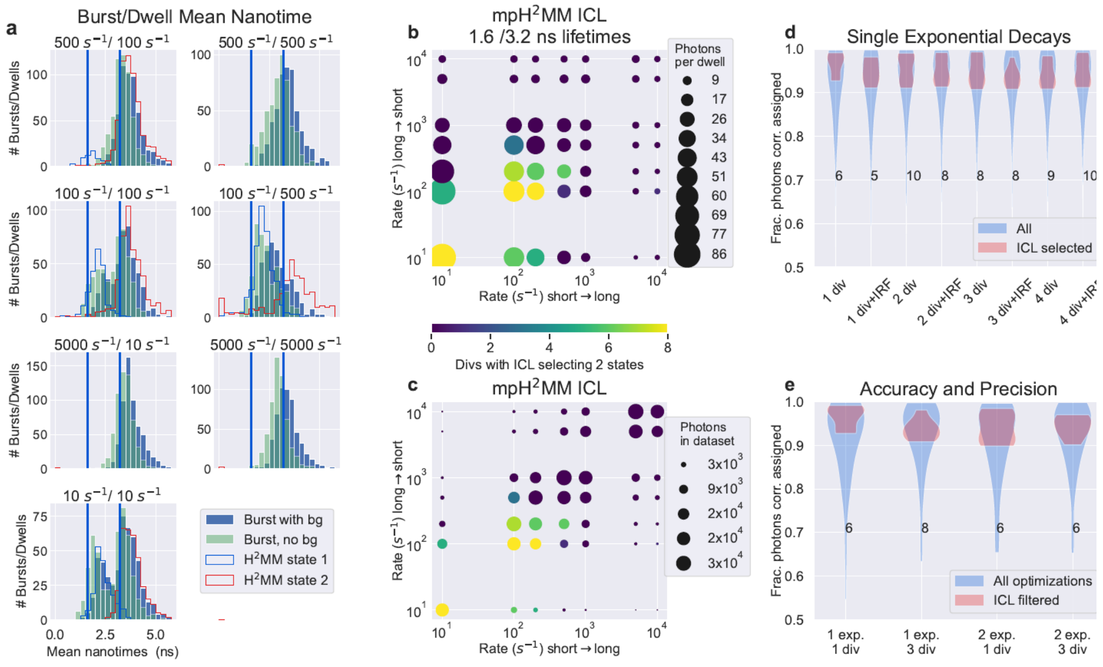

Figure S9. Model selection and accuracy for truncated simulations with states with 1.6 and 3.2 ns lifetimes. Panel arrangement is the same is in Figure 2 of the main text

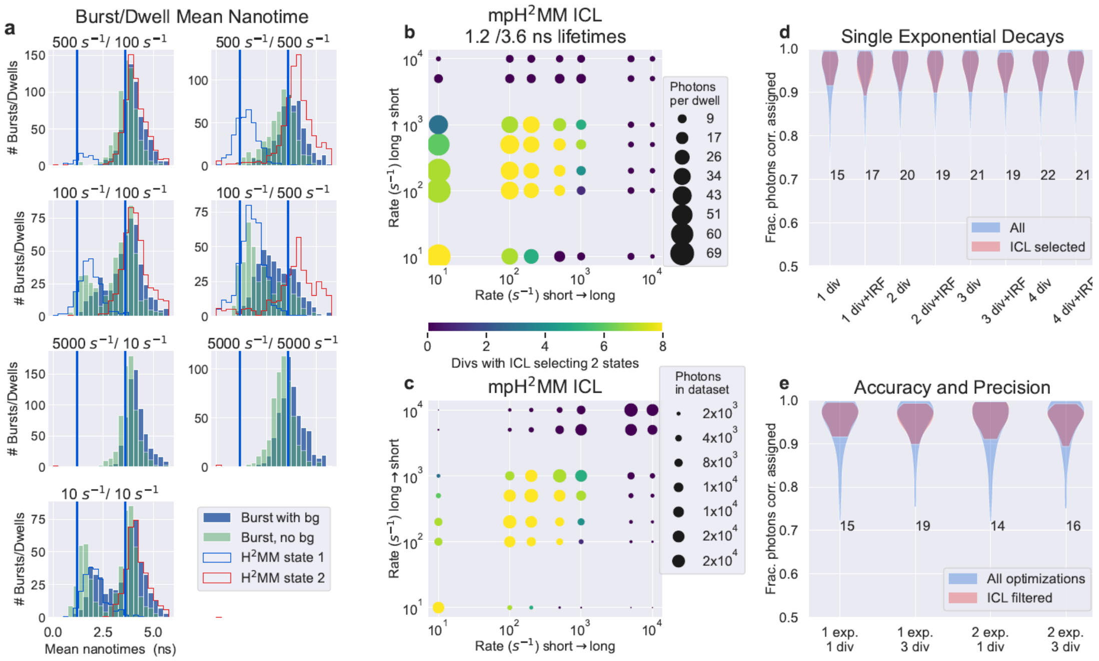

Figure S10. Model selection and accuracy for truncated simulations with states with 1.2 and 3.6 ns lifetimes. Panel arrangement is the same is in Figure 2 of the main text

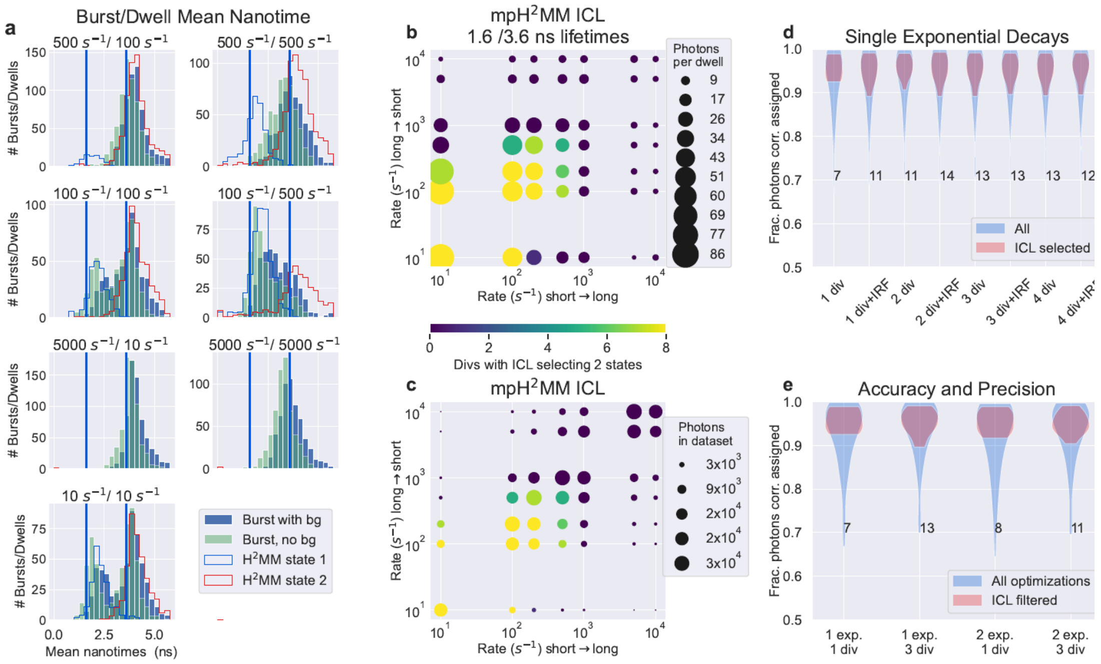

Figure S11. Model selection and accuracy for truncated simulations with states with 1.6 and 3.6 ns lifetimes. Panel arrangement is the same is in Figure 2 of the main text

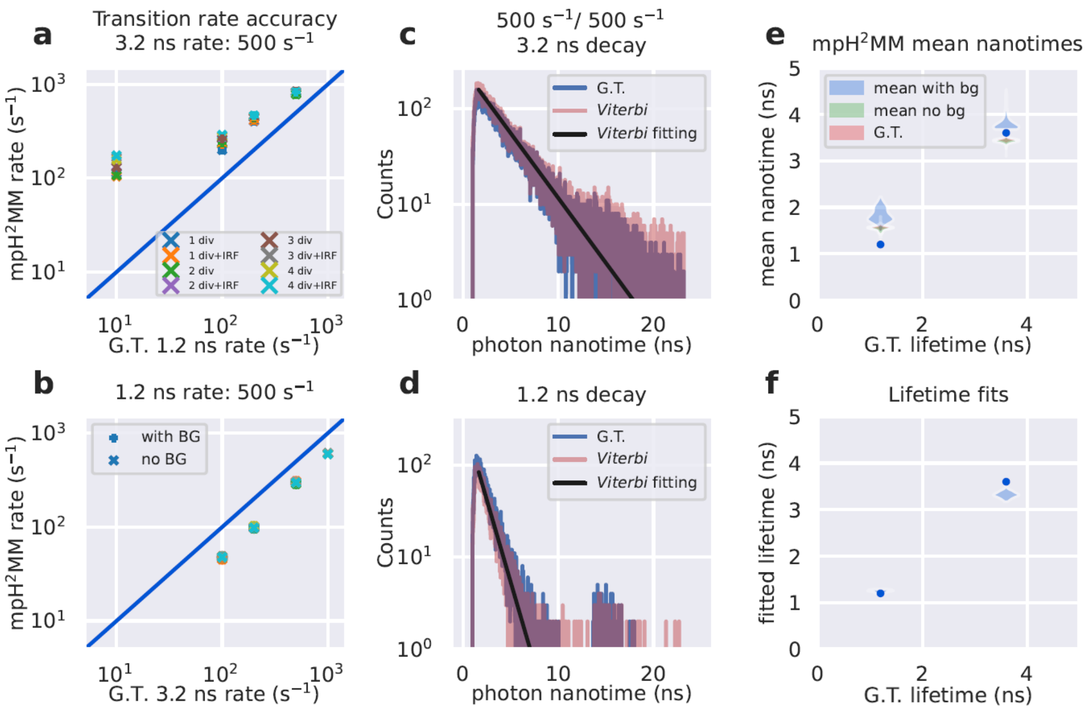

Figure S12. Transition rate and lifetime of ground-truth versus mpH²MM recovered values for simulations of two states with fluorescence lifetimes of 1.2 and 3.2 ns. Panels are arranged the same as in Fig. 3 of the main text. G.T. stands for ground truth.

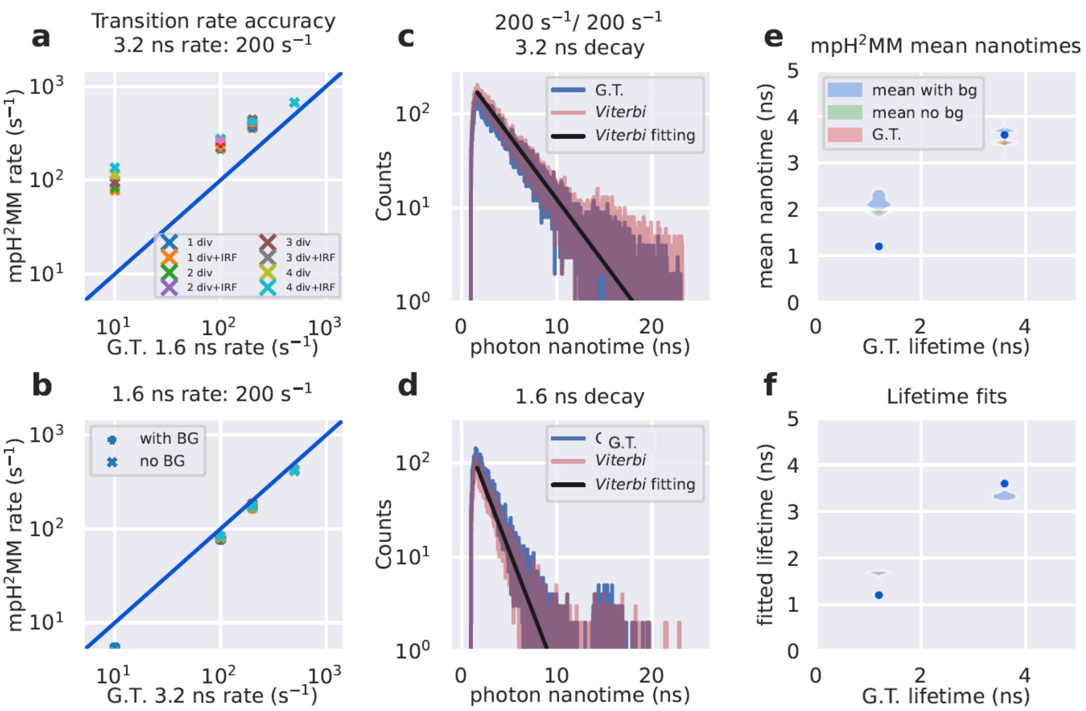

Figure S13. Transition rate and lifetime of ground-truth versus mpH$^2$MM recovered values for simulations of two states with fluorescence lifetimes of 1.6 and 3.2 ns. Panels are arranged the same as in Fig. 3 of the main text. G.T. stands for ground truth.

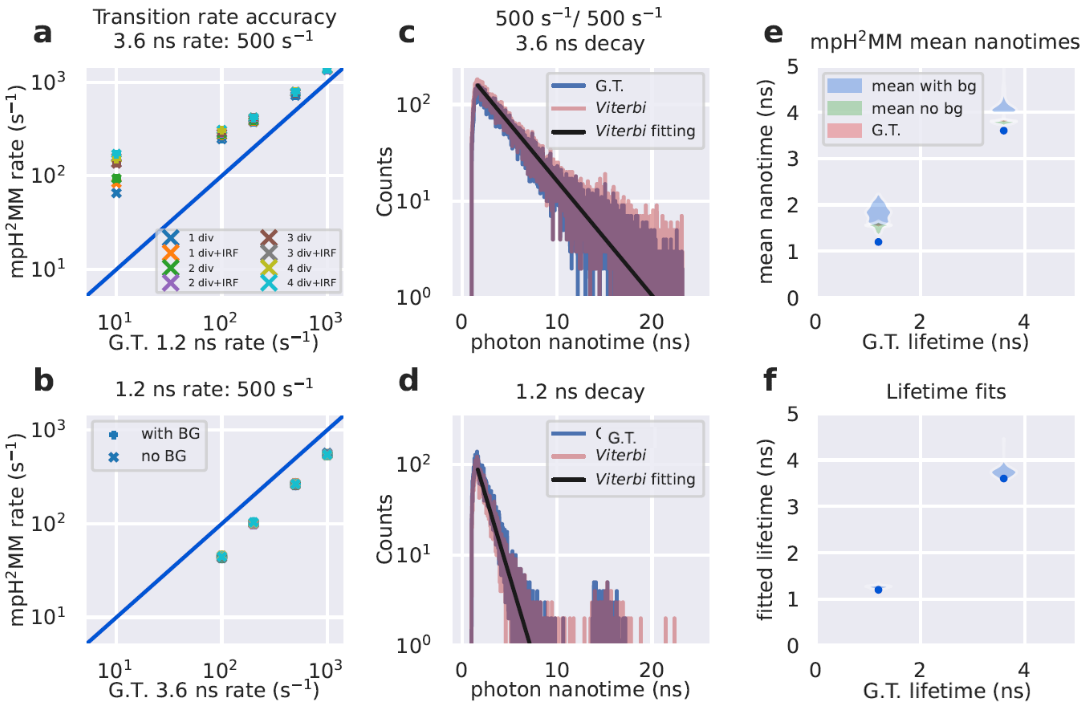

Figure 14. Transition rate and lifetime of ground-truth versus mpH$^2$MM recovered values for simulations of two states with fluorescence lifetimes of 1.2 and 3.6 ns. Panels are arranged the same as in Fig. 3 of the main text. G.T. stands for ground truth.

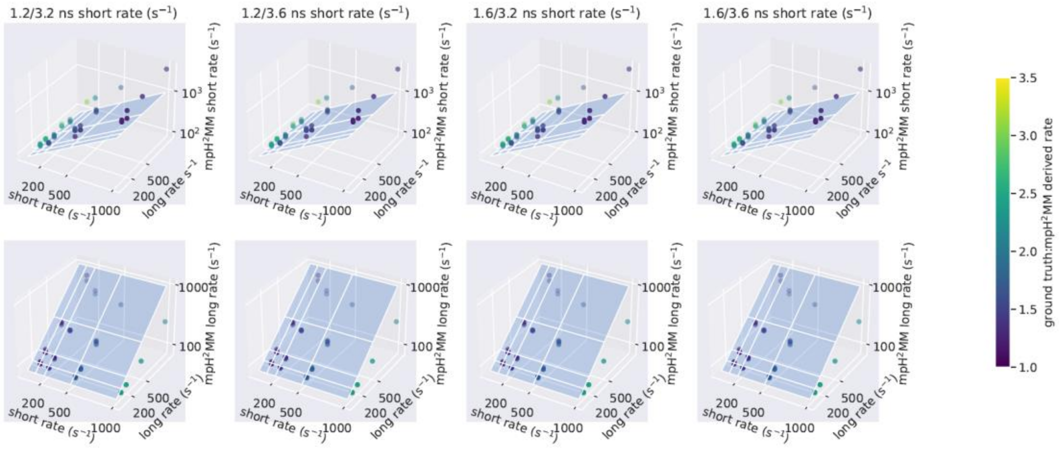

Figure S15. Comparison of ground-truth and recovered transition rates. The ground-truth transition rates are represented on the x and y axes, while the recovered transition rates are represented on the z axis, where the transition rates towards the short or long lifetime states are shown in the upper or lower panels, respectively.

Table S1. Minimum ICL-based selected state models for all simulations and divisor schemes. Ground-truth is always Two-state. Blue indicates ICL selected an under-fit one-state model, green the correct two-state model, and orange an over-fit three-state model. Transition rate values in columns are the transition rates from the long lifetime state to the short lifetime state, while the transition rate values in the upper left above the divisor names are for the short to long lifetime transition rate.

| 1.2 ns / 3.2 ns | | | | | | | |
|---|---|---|---|---|---|---|---|
| [a]1x10$^{1}$ s$^{-1}$ | 1x10$^{1}$ s$^{-1}$ | 1x10$^{2}$ s$^{-1}$ | 2x10$^{2}$ s$^{-1}$ | 5x10$^{2}$ s$^{-1}$ | 1x10$^{3}$ s$^{-1}$ | 5x10$^{3}$ s$^{-1}$ | 1x10$^{4}$ s$^{-1}$ |
| [b]1 div | 2 | 2 | 2 | 1 | 1 | 1 | 1 |
| [c]1 div+IRF | 2 | 3 | 2 | 1 | 1 | 1 | 1 |
| [d]2 div | 2 | 2 | 2 | 1 | 1 | 1 | 1 |
| [e]2 div+IRF | 2 | 2 | 2 | 1 | 1 | 1 | 1 |
| [f]3 div | 2 | 2 | 2 | 1 | 1 | 1 | 1 |
| [g]3 div+IRF | 2 | 2 | 2 | 1 | 1 | 1 | 1 |
| [h]4 div | 2 | 2 | 2 | 1 | 1 | 1 | 1 |
| [i]4 div+IRF | 3 | 2 | 2 | 1 | 1 | 1 | 1 |
| 1x10$^{2}$ s$^{-1}$ | 1x10$^{1}$ s$^{-1}$ | 1x10$^{2}$ s$^{-1}$ | 2x10$^{2}$ s$^{-1}$ | 5x10$^{2}$ s$^{-1}$ | 1x10$^{3}$ s$^{-1}$ | 5x10$^{3}$ s$^{-1}$ | 1x10$^{4}$ s$^{-1}$ |
| 1 div | 2 | 2 | 2 | 2 | 1 | 1 | 1 |
| 1 div+IRF | 1 | 2 | 2 | 2 | 1 | 1 | 1 |
| 2 div | 2 | 2 | 2 | 2 | 1 | 1 | 1 |
| 2 div+IRF | 2 | 2 | 2 | 2 | 2 | 1 | 1 |
| 3 div | 2 | 2 | 2 | 2 | 2 | 1 | 1 |
| 3 div+IRF | 1 | 2 | 2 | 3 | 2 | 1 | 1 |
| 4 div | 2 | 2 | 2 | 3 | 2 | 1 | 1 |
| 4 div+IRF | 2 | 2 | 2 | 3 | 2 | 1 | 1 |
| 2x10$^{2}$ s$^{-1}$ | 1x10$^{1}$ s$^{-1}$ | 1x10$^{2}$ s$^{-1}$ | 2x10$^{2}$ s$^{-1}$ | 5x10$^{2}$ s$^{-1}$ | 1x10$^{3}$ s$^{-1}$ | 5x10$^{3}$ s$^{-1}$ | 1x10$^{4}$ s$^{-1}$ |
| 1 div | 2 | 2 | 2 | 2 | 1 | 1 | 1 |
| 1 div+IRF | 1 | 2 | 2 | 2 | 2 | 1 | 1 |
| 2 div | 2 | 2 | 2 | 2 | 2 | 1 | 1 |
| 2 div+IRF | 2 | 2 | 2 | 2 | 2 | 1 | 1 |
| 3 div | 2 | 2 | 2 | 2 | 2 | 1 | 1 |
| 3 div+IRF | 1 | 2 | 2 | 2 | 2 | 1 | 1 |
| 4 div | 2 | 2 | 2 | 2 | 2 | 1 | 1 |
| 4 div+IRF | 2 | 2 | 2 | 2 | 2 | 1 | 1 |
| 5x10$^{2}$ s$^{-1}$ | 1x10$^{1}$ s$^{-1}$ | 1x10$^{2}$ s$^{-1}$ | 2x10$^{2}$ s$^{-1}$ | 5x10$^{2}$ s$^{-1}$ | 1x10$^{3}$ s$^{-1}$ | 5x10$^{3}$ s$^{-1}$ | 1x10$^{4}$ s$^{-1}$ |

| | | | | | | | |
|---|---|---|---|---|---|---|---|
| 1 div | 1 | 2 | 2 | 1 | 1 | 1 | 1 |
| 1 div+IRF | 1 | 2 | 2 | 2 | 1 | 1 | 1 |
| 2 div | 1 | 2 | 2 | 2 | 1 | 1 | 1 |
| 2 div+IRF | 1 | 2 | 2 | 2 | 1 | 1 | 1 |
| 3 div | 1 | 2 | 2 | 2 | 1 | 1 | 1 |
| 3 div+IRF | 1 | 2 | 2 | 2 | 1 | 1 | 1 |
| 4 div | 1 | 2 | 2 | 2 | 1 | 1 | 1 |
| 4 div+IRF | 1 | 2 | 2 | 2 | 1 | 1 | 1 |
| 1x10$^{3}$ s$^{-1}$ | 1x10$^{1}$ s$^{-1}$ | 1x10$^{2}$ s$^{-1}$ | 2x10$^{2}$ s$^{-1}$ | 5x10$^{2}$ s$^{-1}$ | 1x10$^{3}$ s$^{-1}$ | 5x10$^{3}$ s$^{-1}$ | 1x10$^{4}$ s$^{-1}$ |
| 1 div | 1 | 1 | 1 | 1 | 1 | 1 | 1 |
| 1 div+IRF | 1 | 1 | 2 | 1 | 1 | 1 | 1 |
| 2 div | 1 | 1 | 2 | 1 | 1 | 1 | 1 |
| 2 div+IRF | 1 | 2 | 2 | 1 | 1 | 1 | 1 |
| 3 div | 1 | 2 | 2 | 1 | 1 | 1 | 1 |
| 3 div+IRF | 1 | 1 | 2 | 1 | 1 | 1 | 1 |
| 4 div | 1 | 2 | 2 | 1 | 1 | 1 | 1 |
| 4 div+IRF | 1 | 2 | 2 | 1 | 1 | 1 | 1 |
| 5x10$^{3}$ s$^{-1}$ | 1x10$^{1}$ s$^{-1}$ | 1x10$^{2}$ s$^{-1}$ | 2x10$^{2}$ s$^{-1}$ | 5x10$^{2}$ s$^{-1}$ | 1x10$^{3}$ s$^{-1}$ | 5x10$^{3}$ s$^{-1}$ | 1x10$^{4}$ s$^{-1}$ |
| 1 div | 1 | 1 | 1 | 1 | 1 | 1 | 1 |
| 1 div+IRF | 1 | 1 | 1 | 1 | 1 | 1 | 1 |
| 2 div | 1 | 1 | 1 | 1 | 1 | 1 | 1 |
| 2 div+IRF | 1 | 1 | 1 | 1 | 1 | 1 | 1 |
| 3 div | 1 | 1 | 1 | 1 | 1 | 1 | 1 |
| 3 div+IRF | 1 | 1 | 1 | 1 | 1 | 1 | 1 |
| 4 div | 1 | 1 | 1 | 1 | 1 | 1 | 1 |
| 4 div+IRF | 1 | 1 | 1 | 1 | 1 | 1 | 1 |
| 1x10$^{4}$ s$^{-1}$ | 1x10$^{1}$ s$^{-1}$ | 1x10$^{2}$ s$^{-1}$ | 2x10$^{2}$ s$^{-1}$ | 5x10$^{2}$ s$^{-1}$ | 1x10$^{3}$ s$^{-1}$ | 5x10$^{3}$ s$^{-1}$ | 1x10$^{4}$ s$^{-1}$ |
| 1 div | 1 | 1 | 1 | 1 | 1 | 1 | 1 |
| 1 div+IRF | 1 | 1 | 1 | 1 | 1 | 1 | 1 |
| 2 div | 1 | 1 | 1 | 1 | 1 | 1 | 1 |
| 2 div+IRF | 1 | 1 | 1 | 1 | 1 | 1 | 1 |
| 3 div | 1 | 1 | 1 | 1 | 1 | 1 | 1 |
| 3 div+IRF | 1 | 1 | 1 | 1 | 1 | 1 | 1 |
| 4 div | 1 | 1 | 1 | 1 | 1 | 1 | 1 |
| 4 div+IRF | 1 | 1 | 1 | 1 | 1 | 1 | 1 |
| 1.2 ns / 3.6 ns | | | | | | | |
| 1x10$^{1}$ s$^{-1}$ | 1x10$^{1}$ s$^{-1}$ | 1x10$^{2}$ s$^{-1}$ | 2x10$^{2}$ s$^{-1}$ | 5x10$^{2}$ s$^{-1}$ | 1x10$^{3}$ s$^{-1}$ | 5x10$^{3}$ s$^{-1}$ | 1x10$^{4}$ s$^{-1}$ |
| 1 div | 2 | 2 | 2 | 1 | 1 | 1 | 1 |

| | | | | | | | |
|---|---|---|---|---|---|---|---|
| 1 div+IRF | 2 | 2 | 2 | 1 | 1 | 1 | 1 |
| 2 div | 2 | 3 | 2 | 2 | 1 | 1 | 1 |
| 2 div+IRF | 2 | 3 | 2 | 2 | 1 | 1 | 1 |
| 3 div | 2 | 3 | 2 | 2 | 1 | 1 | 1 |
| 3 div+IRF | 2 | 2 | 2 | 2 | 1 | 1 | 1 |
| 4 div | 2 | 2 | 2 | 2 | 1 | 1 | 1 |
| 4 div+IRF | 2 | 3 | 2 | 2 | 1 | 1 | 1 |
| $1x10^2$ $s^{-1}$ | $1x10^1$ $s^{-1}$ | $1x10^2$ $s^{-1}$ | $2x10^2$ $s^{-1}$ | $5x10^2$ $s^{-1}$ | $1x10^3$ $s^{-1}$ | $5x10^3$ $s^{-1}$ | $1x10^4$ $s^{-1}$ |
| 1 div | 2 | 2 | 2 | 2 | 1 | 1 | 1 |
| 1 div+IRF | 2 | 2 | 2 | 2 | 2 | 1 | 1 |
| 2 div | 2 | 2 | 2 | 2 | 2 | 1 | 1 |
| 2 div+IRF | 2 | 2 | 2 | 2 | 2 | 1 | 1 |
| 3 div | 2 | 2 | 2 | 2 | 2 | 1 | 1 |
| 3 div+IRF | 2 | 2 | 2 | 2 | 2 | 1 | 1 |
| 4 div | 2 | 2 | 2 | 2 | 2 | 1 | 1 |
| 4 div+IRF | 2 | 2 | 2 | 3 | 2 | 1 | 1 |
| $2x10^2$ $s^{-1}$ | $1x10^1$ $s^{-1}$ | $1x10^2$ $s^{-1}$ | $2x10^2$ $s^{-1}$ | $5x10^2$ $s^{-1}$ | $1x10^3$ $s^{-1}$ | $5x10^3$ $s^{-1}$ | $1x10^4$ $s^{-1}$ |
| 1 div | 2 | 2 | 2 | 2 | 2 | 1 | 1 |
| 1 div+IRF | 1 | 2 | 2 | 2 | 2 | 1 | 1 |
| 2 div | 2 | 2 | 2 | 2 | 2 | 1 | 1 |
| 2 div+IRF | 1 | 2 | 2 | 2 | 2 | 1 | 1 |
| 3 div | 1 | 2 | 2 | 2 | 2 | 1 | 1 |
| 3 div+IRF | 1 | 2 | 2 | 2 | 2 | 1 | 1 |
| 4 div | 1 | 2 | 2 | 2 | 2 | 1 | 1 |
| 4 div+IRF | 1 | 2 | 2 | 2 | 2 | 1 | 1 |
| $5x10^2$ $s^{-1}$ | $1x10^1$ $s^{-1}$ | $1x10^2$ $s^{-1}$ | $2x10^2$ $s^{-1}$ | $5x10^2$ $s^{-1}$ | $1x10^3$ $s^{-1}$ | $5x10^3$ $s^{-1}$ | $1x10^4$ $s^{-1}$ |
| 1 div | 1 | 2 | 2 | 2 | 1 | 1 | 1 |
| 1 div+IRF | 1 | 2 | 2 | 2 | 2 | 1 | 1 |
| 2 div | 1 | 2 | 2 | 2 | 2 | 1 | 1 |
| 2 div+IRF | 1 | 2 | 2 | 2 | 2 | 1 | 1 |
| 3 div | 1 | 2 | 2 | 2 | 2 | 1 | 1 |
| 3 div+IRF | 1 | 2 | 2 | 2 | 2 | 1 | 1 |
| 4 div | 1 | 2 | 2 | 2 | 2 | 1 | 1 |
| 4 div+IRF | 1 | 2 | 2 | 2 | 2 | 1 | 1 |
| $1x10^3$ $s^{-1}$ | $1x10^1$ $s^{-1}$ | $1x10^2$ $s^{-1}$ | $2x10^2$ $s^{-1}$ | $5x10^2$ $s^{-1}$ | $1x10^3$ $s^{-1}$ | $5x10^3$ $s^{-1}$ | $1x10^4$ $s^{-1}$ |
| 1 div | 1 | 1 | 1 | 1 | 1 | 1 | 1 |
| 1 div+IRF | 1 | 2 | 2 | 2 | 1 | 1 | 1 |
| 2 div | 1 | 2 | 2 | 2 | 1 | 1 | 1 |
| 2 div+IRF | 1 | 2 | 2 | 2 | 2 | 1 | 1 |

| | | | | | | | |
|---|---|---|---|---|---|---|---|
| 3 div | 1 | 2 | 2 | 2 | 2 | 1 | 1 |
| 3 div+IRF | 1 | 1 | 2 | 2 | 2 | 1 | 1 |
| 4 div | 1 | 2 | 2 | 2 | 2 | 1 | 1 |
| 4 div+IRF | 1 | 2 | 2 | 2 | 2 | 1 | 1 |
| $5x10^3$ $s^{-1}$ | $1x10^1$ $s^{-1}$ | $1x10^2$ $s^{-1}$ | $2x10^2$ $s^{-1}$ | $5x10^2$ $s^{-1}$ | $1x10^3$ $s^{-1}$ | $5x10^3$ $s^{-1}$ | $1x10^4$ $s^{-1}$ |
| 1 div | 1 | 1 | 1 | 1 | 1 | 1 | 1 |
| 1 div+IRF | 1 | 1 | 1 | 1 | 1 | 1 | 1 |
| 2 div | 1 | 1 | 1 | 1 | 1 | 1 | 1 |
| 2 div+IRF | 1 | 1 | 1 | 1 | 1 | 1 | 1 |
| 3 div | 1 | 1 | 1 | 1 | 1 | 1 | 1 |
| 3 div+IRF | 1 | 1 | 1 | 1 | 1 | 1 | 1 |
| 4 div | 1 | 1 | 1 | 1 | 1 | 1 | 1 |
| 4 div+IRF | 1 | 1 | 1 | 1 | 1 | 1 | 1 |
| $1x10^4$ $s^{-1}$ | $1x10^1$ $s^{-1}$ | $1x10^2$ $s^{-1}$ | $2x10^2$ $s^{-1}$ | $5x10^2$ $s^{-1}$ | $1x10^3$ $s^{-1}$ | $5x10^3$ $s^{-1}$ | $1x10^4$ $s^{-1}$ |
| 1 div | 1 | 1 | 1 | 1 | 1 | 1 | 1 |
| 1 div+IRF | 1 | 1 | 1 | 1 | 1 | 1 | 1 |
| 2 div | 1 | 1 | 1 | 1 | 1 | 1 | 1 |
| 2 div+IRF | 1 | 1 | 1 | 1 | 1 | 1 | 1 |
| 3 div | 1 | 1 | 1 | 1 | 1 | 1 | 1 |
| 3 div+IRF | 1 | 1 | 1 | 1 | 1 | 1 | 1 |
| 4 div | 1 | 1 | 1 | 1 | 1 | 1 | 1 |
| 4 div+IRF | 1 | 1 | 1 | 1 | 1 | 1 | 1 |
| 1.6 ns / 3.2 ns | | | | | | | |
| $1x10^1$ $s^{-1}$ | $1x10^1$ $s^{-1}$ | $1x10^2$ $s^{-1}$ | $2x10^2$ $s^{-1}$ | $5x10^2$ $s^{-1}$ | $1x10^3$ $s^{-1}$ | $5x10^3$ $s^{-1}$ | $1x10^4$ $s^{-1}$ |
| 1 div | 2 | 2 | 1 | 1 | 1 | 1 | 1 |
| 1 div+IRF | 2 | 2 | 1 | 1 | 1 | 1 | 1 |
| 2 div | 2 | 2 | 1 | 1 | 1 | 1 | 1 |
| 2 div+IRF | 2 | 2 | 1 | 1 | 1 | 1 | 1 |
| 3 div | 2 | 2 | 1 | 1 | 1 | 1 | 1 |
| 3 div+IRF | 2 | 2 | 1 | 1 | 1 | 1 | 1 |
| 4 div | 2 | 2 | 1 | 1 | 1 | 1 | 1 |
| 4 div+IRF | 2 | 2 | 1 | 1 | 1 | 1 | 1 |
| $1x10^2$ $s^{-1}$ | $1x10^1$ $s^{-1}$ | $1x10^2$ $s^{-1}$ | $2x10^2$ $s^{-1}$ | $5x10^2$ $s^{-1}$ | $1x10^3$ $s^{-1}$ | $5x10^3$ $s^{-1}$ | $1x10^4$ $s^{-1}$ |
| 1 div | 2 | 2 | 1 | 1 | 1 | 1 | 1 |
| 1 div+IRF | 1 | 2 | 2 | 1 | 1 | 1 | 1 |
| 2 div | 2 | 2 | 2 | 1 | 1 | 1 | 1 |
| 2 div+IRF | 2 | 2 | 2 | 1 | 1 | 1 | 1 |
| 3 div | 2 | 2 | 2 | 1 | 1 | 1 | 1 |

| | | | | | | | |
|---|---|---|---|---|---|---|---|
| 3 div+IRF | 1 | 2 | 2 | 1 | 1 | 1 | 1 |
| 4 div | 2 | 2 | 3 | 1 | 1 | 1 | 1 |
| 4 div+IRF | 2 | 2 | 3 | 1 | 1 | 1 | 1 |
| 2x10$^2$ s$^{-1}$ | 1x10$^1$ s$^{-1}$ | 1x10$^2$ s$^{-1}$ | 2x10$^2$ s$^{-1}$ | 5x10$^2$ s$^{-1}$ | 1x10$^3$ s$^{-1}$ | 5x10$^3$ s$^{-1}$ | 1x10$^4$ s$^{-1}$ |
| 1 div | 2 | 2 | 1 | 1 | 1 | 1 | 1 |
| 1 div+IRF | 1 | 2 | 2 | 1 | 1 | 1 | 1 |
| 2 div | 2 | 2 | 2 | 1 | 1 | 1 | 1 |
| 2 div+IRF | 2 | 2 | 2 | 1 | 1 | 1 | 1 |
| 3 div | 1 | 2 | 2 | 1 | 1 | 1 | 1 |
| 3 div+IRF | 1 | 2 | 2 | 1 | 1 | 1 | 1 |
| 4 div | 1 | 2 | 2 | 1 | 1 | 1 | 1 |
| 4 div+IRF | 2 | 2 | 2 | 1 | 1 | 1 | 1 |
| 5x10$^2$ s$^{-1}$ | 1x10$^1$ s$^{-1}$ | 1x10$^2$ s$^{-1}$ | 2x10$^2$ s$^{-1}$ | 5x10$^2$ s$^{-1}$ | 1x10$^3$ s$^{-1}$ | 5x10$^3$ s$^{-1}$ | 1x10$^4$ s$^{-1}$ |
| 1 div | 1 | 1 | 1 | 1 | 1 | 1 | 1 |
| 1 div+IRF | 1 | 1 | 1 | 1 | 1 | 1 | 1 |
| 2 div | 1 | 1 | 1 | 1 | 1 | 1 | 1 |
| 2 div+IRF | 1 | 1 | 1 | 1 | 1 | 1 | 1 |
| 3 div | 1 | 1 | 1 | 1 | 1 | 1 | 1 |
| 3 div+IRF | 1 | 2 | 1 | 1 | 1 | 1 | 1 |
| 4 div | 1 | 1 | 1 | 1 | 1 | 1 | 1 |
| 4 div+IRF | 1 | 2 | 1 | 1 | 1 | 1 | 1 |
| 1x10$^3$ s$^{-1}$ | 1x10$^1$ s$^{-1}$ | 1x10$^2$ s$^{-1}$ | 2x10$^2$ s$^{-1}$ | 5x10$^2$ s$^{-1}$ | 1x10$^3$ s$^{-1}$ | 5x10$^3$ s$^{-1}$ | 1x10$^4$ s$^{-1}$ |
| 1 div | 1 | 1 | 1 | 1 | 1 | 1 | 1 |
| 1 div+IRF | 1 | 1 | 1 | 1 | 1 | 1 | 1 |
| 2 div | 1 | 1 | 1 | 1 | 1 | 1 | 1 |
| 2 div+IRF | 1 | 1 | 1 | 1 | 1 | 1 | 1 |
| 3 div | 1 | 1 | 1 | 1 | 1 | 1 | 1 |
| 3 div+IRF | 1 | 1 | 1 | 1 | 1 | 1 | 1 |
| 4 div | 1 | 1 | 1 | 1 | 1 | 1 | 1 |
| 4 div+IRF | 1 | 1 | 1 | 1 | 1 | 1 | 1 |
| 5x10$^3$ s$^{-1}$ | 1x10$^1$ s$^{-1}$ | 1x10$^2$ s$^{-1}$ | 2x10$^2$ s$^{-1}$ | 5x10$^2$ s$^{-1}$ | 1x10$^3$ s$^{-1}$ | 5x10$^3$ s$^{-1}$ | 1x10$^4$ s$^{-1}$ |
| 1 div | 1 | 1 | 1 | 1 | 1 | 1 | 1 |
| 1 div+IRF | 1 | 1 | 1 | 1 | 1 | 1 | 1 |
| 2 div | 1 | 1 | 1 | 1 | 1 | 1 | 1 |
| 2 div+IRF | 1 | 1 | 1 | 1 | 1 | 1 | 1 |
| 3 div | 1 | 1 | 1 | 1 | 1 | 1 | 1 |
| 3 div+IRF | 1 | 1 | 1 | 1 | 1 | 1 | 1 |
| 4 div | 1 | 1 | 1 | 1 | 1 | 1 | 1 |
| 4 div+IRF | 1 | 1 | 1 | 1 | 1 | 1 | 1 |

| | | | | | | | |
|---|---|---|---|---|---|---|---|
| 1x10$^{4}$ s$^{-1}$ | 1x10$^{1}$ s$^{-1}$ | 1x10$^{2}$ s$^{-1}$ | 2x10$^{2}$ s$^{-1}$ | 5x10$^{2}$ s$^{-1}$ | 1x10$^{3}$ s$^{-1}$ | 5x10$^{3}$ s$^{-1}$ | 1x10$^{4}$ s$^{-1}$ |
| 1 div | 1 | 1 | 1 | 1 | 1 | 1 | 1 |
| 1 div+IRF | 1 | 1 | 1 | 1 | 1 | 1 | 1 |
| 2 div | 1 | 1 | 1 | 1 | 1 | 1 | 1 |
| 2 div+IRF | 1 | 1 | 1 | 1 | 1 | 1 | 1 |
| 3 div | 1 | 1 | 1 | 1 | 1 | 1 | 1 |
| 3 div+IRF | 1 | 1 | 1 | 1 | 1 | 1 | 1 |
| 4 div | 1 | 1 | 1 | 1 | 1 | 1 | 1 |
| 4 div+IRF | 1 | 1 | 1 | 1 | 1 | 1 | 1 |
| 1.6 ns / 3.6 ns | | | | | | | |
| 1x10$^{1}$ s$^{-1}$ | 1x10$^{1}$ s$^{-1}$ | 1x10$^{2}$ s$^{-1}$ | 2x10$^{2}$ s$^{-1}$ | 5x10$^{2}$ s$^{-1}$ | 1x10$^{3}$ s$^{-1}$ | 5x10$^{3}$ s$^{-1}$ | 1x10$^{4}$ s$^{-1}$ |
| 1 div | 2 | 2 | 1 | 1 | 1 | 1 | 1 |
| 1 div+IRF | 2 | 2 | 1 | 1 | 1 | 1 | 1 |
| 2 div | 2 | 3 | 1 | 1 | 1 | 1 | 1 |
| 2 div+IRF | 2 | 3 | 2 | 1 | 1 | 1 | 1 |
| 3 div | 2 | 3 | 2 | 1 | 1 | 1 | 1 |
| 3 div+IRF | 2 | 3 | 3 | 1 | 1 | 1 | 1 |
| 4 div | 2 | 3 | 2 | 1 | 1 | 1 | 1 |
| 4 div+IRF | 2 | 2 | 2 | 1 | 1 | 1 | 1 |
| 1x10$^{2}$ s$^{-1}$ | 1x10$^{1}$ s$^{-1}$ | 1x10$^{2}$ s$^{-1}$ | 2x10$^{2}$ s$^{-1}$ | 5x10$^{2}$ s$^{-1}$ | 1x10$^{3}$ s$^{-1}$ | 5x10$^{3}$ s$^{-1}$ | 1x10$^{4}$ s$^{-1}$ |
| 1 div | 2 | 2 | 2 | 1 | 1 | 1 | 1 |
| 1 div+IRF | 2 | 2 | 2 | 2 | 1 | 1 | 1 |
| 2 div | 2 | 2 | 2 | 2 | 1 | 1 | 1 |
| 2 div+IRF | 2 | 2 | 2 | 2 | 1 | 1 | 1 |
| 3 div | 2 | 2 | 2 | 2 | 1 | 1 | 1 |
| 3 div+IRF | 2 | 2 | 2 | 2 | 1 | 1 | 1 |
| 4 div | 2 | 2 | 2 | 2 | 1 | 1 | 1 |
| 4 div+IRF | 2 | 2 | 2 | 2 | 1 | 1 | 1 |
| 2x10$^{2}$ s$^{-1}$ | 1x10$^{1}$ s$^{-1}$ | 1x10$^{2}$ s$^{-1}$ | 2x10$^{2}$ s$^{-1}$ | 5x10$^{2}$ s$^{-1}$ | 1x10$^{3}$ s$^{-1}$ | 5x10$^{3}$ s$^{-1}$ | 1x10$^{4}$ s$^{-1}$ |
| 1 div | 1 | 2 | 2 | 1 | 1 | 1 | 1 |
| 1 div+IRF | 1 | 2 | 2 | 2 | 1 | 1 | 1 |
| 2 div | 1 | 2 | 2 | 2 | 1 | 1 | 1 |
| 2 div+IRF | 2 | 2 | 2 | 2 | 1 | 1 | 1 |
| 3 div | 1 | 2 | 2 | 2 | 1 | 1 | 1 |
| 3 div+IRF | 1 | 2 | 2 | 2 | 1 | 1 | 1 |
| 4 div | 1 | 2 | 2 | 2 | 1 | 1 | 1 |
| 4 div+IRF | 1 | 2 | 2 | 2 | 1 | 1 | 1 |

| 5x10$^{2}$ s$^{-1}$ | 1x10$^{1}$ s$^{-1}$ | 1x10$^{2}$ s$^{-1}$ | 2x10$^{2}$ s$^{-1}$ | 5x10$^{2}$ s$^{-1}$ | 1x10$^{3}$ s$^{-1}$ | 5x10$^{3}$ s$^{-1}$ | 1x10$^{4}$ s$^{-1}$ |
|---|---|---|---|---|---|---|---|
| 1 div | 1 | 1 | 1 | 1 | 1 | 1 | 1 |
| 1 div+IRF | 1 | 2 | 2 | 1 | 1 | 1 | 1 |
| 2 div | 1 | 2 | 2 | 1 | 1 | 1 | 1 |
| 2 div+IRF | 1 | 2 | 2 | 2 | 1 | 1 | 1 |
| 3 div | 1 | 2 | 2 | 2 | 1 | 1 | 1 |
| 3 div+IRF | 1 | 2 | 2 | 2 | 1 | 1 | 1 |
| 4 div | 1 | 2 | 2 | 2 | 1 | 1 | 1 |
| 4 div+IRF | 1 | 2 | 2 | 2 | 1 | 1 | 1 |
| 1x10$^{3}$ s$^{-1}$ | 1x10$^{1}$ s$^{-1}$ | 1x10$^{2}$ s$^{-1}$ | 2x10$^{2}$ s$^{-1}$ | 5x10$^{2}$ s$^{-1}$ | 1x10$^{3}$ s$^{-1}$ | 5x10$^{3}$ s$^{-1}$ | 1x10$^{4}$ s$^{-1}$ |
| 1 div | 1 | 1 | 1 | 1 | 1 | 1 | 1 |
| 1 div+IRF | 1 | 1 | 1 | 1 | 1 | 1 | 1 |
| 2 div | 1 | 1 | 1 | 1 | 1 | 1 | 1 |
| 2 div+IRF | 1 | 1 | 1 | 1 | 1 | 1 | 1 |
| 3 div | 1 | 1 | 1 | 1 | 1 | 1 | 1 |
| 3 div+IRF | 1 | 1 | 1 | 1 | 1 | 1 | 1 |
| 4 div | 1 | 1 | 1 | 1 | 1 | 1 | 1 |
| 4 div+IRF | 1 | 1 | 1 | 1 | 1 | 1 | 1 |
| 5x10$^{3}$ s$^{-1}$ | 1x10$^{1}$ s$^{-1}$ | 1x10$^{2}$ s$^{-1}$ | 2x10$^{2}$ s$^{-1}$ | 5x10$^{2}$ s$^{-1}$ | 1x10$^{3}$ s$^{-1}$ | 5x10$^{3}$ s$^{-1}$ | 1x10$^{4}$ s$^{-1}$ |
| 1 div | 1 | 1 | 1 | 1 | 1 | 1 | 1 |
| 1 div+IRF | 1 | 1 | 1 | 1 | 1 | 1 | 1 |
| 2 div | 1 | 1 | 1 | 1 | 1 | 1 | 1 |
| 2 div+IRF | 1 | 1 | 1 | 1 | 1 | 1 | 1 |
| 3 div | 1 | 1 | 1 | 1 | 1 | 1 | 1 |
| 3 div+IRF | 1 | 1 | 1 | 1 | 1 | 1 | 1 |
| 4 div | 1 | 1 | 1 | 1 | 1 | 1 | 1 |
| 4 div+IRF | 1 | 1 | 1 | 1 | 1 | 1 | 1 |
| 1x10$^{4}$ s$^{-1}$ | 1x10$^{1}$ s$^{-1}$ | 1x10$^{2}$ s$^{-1}$ | 2x10$^{2}$ s$^{-1}$ | 5x10$^{2}$ s$^{-1}$ | 1x10$^{3}$ s$^{-1}$ | 5x10$^{3}$ s$^{-1}$ | 1x10$^{4}$ s$^{-1}$ |
| 1 div | 1 | 1 | 1 | 1 | 1 | 1 | 1 |
| 1 div+IRF | 1 | 1 | 1 | 1 | 1 | 1 | 1 |
| 2 div | 1 | 1 | 1 | 1 | 1 | 1 | 1 |
| 2 div+IRF | 1 | 1 | 1 | 1 | 1 | 1 | 1 |
| 3 div | 1 | 1 | 1 | 1 | 1 | 1 | 1 |
| 3 div+IRF | 1 | 1 | 1 | 1 | 1 | 1 | 1 |
| 4 div | 1 | 1 | 1 | 1 | 1 | 1 | 1 |
| 4 div+IRF | 1 | 1 | 1 | 1 | 1 | 1 | 1 |
| multiexponential 1.2 ns / 3.2 ns | | | | | | | |
| 1x10$^{1}$ s$^{-1}$ | 1x10$^{1}$ s$^{-1}$ | 1x10$^{2}$ s$^{-1}$ | 2x10$^{2}$ s$^{-1}$ | 5x10$^{2}$ s$^{-1}$ | 1x10$^{3}$ s$^{-1}$ | 5x10$^{3}$ s$^{-1}$ | 1x10$^{4}$ s$^{-1}$ |

| | | | | | | | |
|---|---|---|---|---|---|---|---|
| 1 div | 2 | 2 | 1 | 1 | 1 | 1 | 1 |
| 1 div+IRF | 2 | 3 | 1 | 1 | 1 | 1 | 1 |
| 2 div | 2 | 3 | 1 | 1 | 1 | 1 | 1 |
| 2 div+IRF | 2 | 2 | 2 | 1 | 1 | 1 | 1 |
| 3 div | 2 | 2 | 2 | 1 | 1 | 1 | 1 |
| 3 div+IRF | 2 | 3 | 2 | 1 | 1 | 1 | 1 |
| 4 div | 2 | 2 | 2 | 1 | 1 | 1 | 1 |
| 4 div+IRF | 2 | 3 | 2 | 1 | 1 | 1 | 1 |
| 1x10$^{2}$ s$^{-1}$ | 1x10$^{1}$ s$^{-1}$ | 1x10$^{2}$ s$^{-1}$ | 2x10$^{2}$ s$^{-1}$ | 5x10$^{2}$ s$^{-1}$ | 1x10$^{3}$ s$^{-1}$ | 5x10$^{3}$ s$^{-1}$ | 1x10$^{4}$ s$^{-1}$ |
| 1 div | 2 | 2 | 2 | 1 | 1 | 1 | 1 |
| 1 div+IRF | 1 | 2 | 2 | 2 | 1 | 1 | 1 |
| 2 div | 2 | 2 | 2 | 2 | 1 | 1 | 1 |
| 2 div+IRF | 2 | 2 | 2 | 2 | 2 | 1 | 1 |
| 3 div | 2 | 2 | 2 | 2 | 2 | 1 | 1 |
| 3 div+IRF | 2 | 2 | 2 | 2 | 2 | 1 | 1 |
| 4 div | 2 | 2 | 2 | 2 | 2 | 1 | 1 |
| 4 div+IRF | 2 | 2 | 3 | 3 | 2 | 1 | 1 |
| 2x10$^{2}$ s$^{-1}$ | 1x10$^{1}$ s$^{-1}$ | 1x10$^{2}$ s$^{-1}$ | 2x10$^{2}$ s$^{-1}$ | 5x10$^{2}$ s$^{-1}$ | 1x10$^{3}$ s$^{-1}$ | 5x10$^{3}$ s$^{-1}$ | 1x10$^{4}$ s$^{-1}$ |
| 1 div | 1 | 2 | 2 | 2 | 1 | 1 | 1 |
| 1 div+IRF | 1 | 2 | 2 | 2 | 1 | 1 | 1 |
| 2 div | 1 | 2 | 2 | 2 | 1 | 1 | 1 |
| 2 div+IRF | 2 | 2 | 2 | 2 | 2 | 1 | 1 |
| 3 div | 1 | 2 | 2 | 2 | 2 | 1 | 1 |
| 3 div+IRF | 1 | 2 | 2 | 2 | 2 | 1 | 1 |
| 4 div | 1 | 2 | 2 | 2 | 2 | 1 | 1 |
| 4 div+IRF | 1 | 2 | 2 | 2 | 2 | 1 | 1 |
| 5x10$^{2}$ s$^{-1}$ | 1x10$^{1}$ s$^{-1}$ | 1x10$^{2}$ s$^{-1}$ | 2x10$^{2}$ s$^{-1}$ | 5x10$^{2}$ s$^{-1}$ | 1x10$^{3}$ s$^{-1}$ | 5x10$^{3}$ s$^{-1}$ | 1x10$^{4}$ s$^{-1}$ |
| 1 div | 1 | 2 | 2 | 1 | 1 | 1 | 1 |
| 1 div+IRF | 1 | 2 | 2 | 1 | 1 | 1 | 1 |
| 2 div | 1 | 2 | 2 | 2 | 1 | 1 | 1 |
| 2 div+IRF | 1 | 2 | 2 | 2 | 1 | 1 | 1 |
| 3 div | 1 | 2 | 2 | 2 | 1 | 1 | 1 |
| 3 div+IRF | 1 | 2 | 2 | 2 | 1 | 1 | 1 |
| 4 div | 1 | 2 | 2 | 2 | 1 | 1 | 1 |
| 4 div+IRF | 1 | 2 | 2 | 2 | 1 | 1 | 1 |
| 1x10$^{3}$ s$^{-1}$ | 1x10$^{1}$ s$^{-1}$ | 1x10$^{2}$ s$^{-1}$ | 2x10$^{2}$ s$^{-1}$ | 5x10$^{2}$ s$^{-1}$ | 1x10$^{3}$ s$^{-1}$ | 5x10$^{3}$ s$^{-1}$ | 1x10$^{4}$ s$^{-1}$ |
| 1 div | 1 | 1 | 1 | 1 | 1 | 1 | 1 |
| 1 div+IRF | 1 | 1 | 1 | 1 | 1 | 1 | 1 |
| 2 div | 1 | 1 | 1 | 1 | 1 | 1 | 1 |

| | | | | | | | |
|---|---|---|---|---|---|---|---|
| 2 div+IRF | 1 | 1 | 2 | 1 | 1 | 1 | 1 |
| 3 div | 1 | 1 | 2 | 1 | 1 | 1 | 1 |
| 3 div+IRF | 1 | 1 | 1 | 1 | 1 | 1 | 1 |
| 4 div | 1 | 1 | 2 | 1 | 1 | 1 | 1 |
| 4 div+IRF | 1 | 1 | 2 | 1 | 1 | 1 | 1 |
| 5x10$^{3}$ s$^{-1}$ | 1x10$^{1}$ s$^{-1}$ | 1x10$^{2}$ s$^{-1}$ | 2x10$^{2}$ s$^{-1}$ | 5x10$^{2}$ s$^{-1}$ | 1x10$^{3}$ s$^{-1}$ | 5x10$^{3}$ s$^{-1}$ | 1x10$^{4}$ s$^{-1}$ |
| 1 div | 1 | 1 | 1 | 1 | 1 | 1 | 1 |
| 1 div+IRF | 1 | 1 | 1 | 1 | 1 | 1 | 1 |
| 2 div | 1 | 1 | 1 | 1 | 1 | 1 | 1 |
| 2 div+IRF | 1 | 1 | 1 | 1 | 1 | 1 | 1 |
| 3 div | 1 | 1 | 1 | 1 | 1 | 1 | 1 |
| 3 div+IRF | 1 | 1 | 1 | 1 | 1 | 1 | 1 |
| 4 div | 1 | 1 | 1 | 1 | 1 | 1 | 1 |
| 4 div+IRF | 1 | 1 | 1 | 1 | 1 | 1 | 1 |
| 1x10$^{4}$ s$^{-1}$ | 1x10$^{1}$ s$^{-1}$ | 1x10$^{2}$ s$^{-1}$ | 2x10$^{2}$ s$^{-1}$ | 5x10$^{2}$ s$^{-1}$ | 1x10$^{3}$ s$^{-1}$ | 5x10$^{3}$ s$^{-1}$ | 1x10$^{4}$ s$^{-1}$ |
| 1 div | 1 | 1 | 1 | 1 | 1 | 1 | 1 |
| 1 div+IRF | 1 | 1 | 1 | 1 | 1 | 1 | 1 |
| 2 div | 1 | 1 | 1 | 1 | 1 | 1 | 1 |
| 2 div+IRF | 1 | 1 | 1 | 1 | 1 | 1 | 1 |
| 3 div | 1 | 1 | 1 | 1 | 1 | 1 | 1 |
| 3 div+IRF | 1 | 1 | 1 | 1 | 1 | 1 | 1 |
| 4 div | 1 | 1 | 1 | 1 | 1 | 1 | 1 |
| 4 div+IRF | 1 | 1 | 1 | 1 | 1 | 1 | 1 |
| multiexponential 1.2 ns / 3.6 ns | | | | | | | |
| 1x10$^{1}$ s$^{-1}$ | 1x10$^{1}$ s$^{-1}$ | 1x10$^{2}$ s$^{-1}$ | 2x10$^{2}$ s$^{-1}$ | 5x10$^{2}$ s$^{-1}$ | 1x10$^{3}$ s$^{-1}$ | 5x10$^{3}$ s$^{-1}$ | 1x10$^{4}$ s$^{-1}$ |
| 1 div | 2 | 2 | 3 | 1 | 1 | 1 | 1 |
| 1 div+IRF | 2 | 3 | 2 | 1 | 1 | 1 | 1 |
| 2 div | 2 | 3 | 2 | 1 | 1 | 1 | 1 |
| 2 div+IRF | 2 | 3 | 2 | 2 | 1 | 1 | 1 |
| 3 div | 2 | 2 | 2 | 2 | 1 | 1 | 1 |
| 3 div+IRF | 2 | 3 | 3 | 2 | 1 | 1 | 1 |
| 4 div | 2 | 3 | 2 | 2 | 1 | 1 | 1 |
| 4 div+IRF | 2 | 3 | 2 | 2 | 1 | 1 | 1 |
| 1x10$^{2}$ s$^{-1}$ | 1x10$^{1}$ s$^{-1}$ | 1x10$^{2}$ s$^{-1}$ | 2x10$^{2}$ s$^{-1}$ | 5x10$^{2}$ s$^{-1}$ | 1x10$^{3}$ s$^{-1}$ | 5x10$^{3}$ s$^{-1}$ | 1x10$^{4}$ s$^{-1}$ |
| 1 div | 2 | 2 | 2 | 2 | 1 | 1 | 1 |
| 1 div+IRF | 1 | 2 | 2 | 2 | 1 | 1 | 1 |
| 2 div | 2 | 2 | 2 | 2 | 1 | 1 | 1 |
| 2 div+IRF | 2 | 2 | 2 | 2 | 2 | 1 | 1 |

| | | | | | | | |
|---|---|---|---|---|---|---|---|
| 3 div | 2 | 2 | 2 | 2 | 2 | 1 | 1 |
| 3 div+IRF | 3 | 2 | 2 | 2 | 2 | 1 | 1 |
| 4 div | 2 | 2 | 2 | 2 | 2 | 1 | 1 |
| 4 div+IRF | 2 | 2 | 2 | 2 | 2 | 1 | 1 |
| 2x10$^{2}$ s$^{-1}$ | 1x10$^{1}$ s$^{-1}$ | 1x10$^{2}$ s$^{-1}$ | 2x10$^{2}$ s$^{-1}$ | 5x10$^{2}$ s$^{-1}$ | 1x10$^{3}$ s$^{-1}$ | 5x10$^{3}$ s$^{-1}$ | 1x10$^{4}$ s$^{-1}$ |
| 1 div | 2 | 2 | 2 | 2 | 1 | 1 | 1 |
| 1 div+IRF | 1 | 2 | 2 | 2 | 2 | 1 | 1 |
| 2 div | 2 | 2 | 2 | 2 | 2 | 1 | 1 |
| 2 div+IRF | 2 | 2 | 2 | 2 | 2 | 1 | 1 |
| 3 div | 2 | 2 | 2 | 2 | 2 | 1 | 1 |
| 3 div+IRF | 1 | 2 | 2 | 2 | 2 | 1 | 1 |
| 4 div | 2 | 2 | 2 | 2 | 2 | 1 | 1 |
| 4 div+IRF | 2 | 2 | 2 | 2 | 2 | 1 | 1 |
| 5x10$^{2}$ s$^{-1}$ | 1x10$^{1}$ s$^{-1}$ | 1x10$^{2}$ s$^{-1}$ | 2x10$^{2}$ s$^{-1}$ | 5x10$^{2}$ s$^{-1}$ | 1x10$^{3}$ s$^{-1}$ | 5x10$^{3}$ s$^{-1}$ | 1x10$^{4}$ s$^{-1}$ |
| 1 div | 1 | 2 | 2 | 2 | 1 | 1 | 1 |
| 1 div+IRF | 1 | 2 | 2 | 2 | 2 | 1 | 1 |
| 2 div | 1 | 2 | 2 | 2 | 2 | 1 | 1 |
| 2 div+IRF | 2 | 2 | 2 | 2 | 2 | 1 | 1 |
| 3 div | 1 | 2 | 2 | 2 | 2 | 1 | 1 |
| 3 div+IRF | 1 | 2 | 2 | 2 | 2 | 1 | 1 |
| 4 div | 1 | 2 | 2 | 2 | 2 | 1 | 1 |
| 4 div+IRF | 1 | 2 | 2 | 2 | 2 | 1 | 1 |
| 1x10$^{3}$ s$^{-1}$ | 1x10$^{1}$ s$^{-1}$ | 1x10$^{2}$ s$^{-1}$ | 2x10$^{2}$ s$^{-1}$ | 5x10$^{2}$ s$^{-1}$ | 1x10$^{3}$ s$^{-1}$ | 5x10$^{3}$ s$^{-1}$ | 1x10$^{4}$ s$^{-1}$ |
| 1 div | 1 | 2 | 1 | 1 | 1 | 1 | 1 |
| 1 div+IRF | 1 | 1 | 1 | 1 | 1 | 1 | 1 |
| 2 div | 1 | 2 | 1 | 1 | 1 | 1 | 1 |
| 2 div+IRF | 1 | 2 | 2 | 1 | 1 | 1 | 1 |
| 3 div | 1 | 2 | 2 | 1 | 1 | 1 | 1 |
| 3 div+IRF | 1 | 1 | 2 | 2 | 1 | 1 | 1 |
| 4 div | 1 | 2 | 2 | 2 | 1 | 1 | 1 |
| 4 div+IRF | 1 | 2 | 2 | 2 | 1 | 1 | 1 |
| 5x10$^{3}$ s$^{-1}$ | 1x10$^{1}$ s$^{-1}$ | 1x10$^{2}$ s$^{-1}$ | 2x10$^{2}$ s$^{-1}$ | 5x10$^{2}$ s$^{-1}$ | 1x10$^{3}$ s$^{-1}$ | 5x10$^{3}$ s$^{-1}$ | 1x10$^{4}$ s$^{-1}$ |
| 1 div | 1 | 1 | 1 | 1 | 1 | 1 | 1 |
| 1 div+IRF | 1 | 1 | 1 | 1 | 1 | 1 | 1 |
| 2 div | 1 | 1 | 1 | 1 | 1 | 1 | 1 |
| 2 div+IRF | 1 | 1 | 1 | 1 | 1 | 1 | 1 |
| 3 div | 1 | 1 | 1 | 1 | 1 | 1 | 1 |
| 3 div+IRF | 1 | 1 | 1 | 1 | 1 | 1 | 1 |
| 4 div | 1 | 1 | 1 | 1 | 1 | 1 | 1 |

| | | | | | | | |
|---|---|---|---|---|---|---|---|
| 4 div+IRF | 1 | 1 | 1 | 1 | 1 | 1 | 1 |
| $1x10^4 s^{-1}$ | $1x10^1 s^{-1}$ | $1x10^2 s^{-1}$ | $2x10^2 s^{-1}$ | $5x10^2 s^{-1}$ | $1x10^3 s^{-1}$ | $5x10^3 s^{-1}$ | $1x10^4 s^{-1}$ |
| 1 div | 1 | 1 | 1 | 1 | 1 | 1 | 1 |
| 1 div+IRF | 1 | 1 | 1 | 1 | 1 | 1 | 1 |
| 2 div | 1 | 1 | 1 | 1 | 1 | 1 | 1 |
| 2 div+IRF | 1 | 1 | 1 | 1 | 1 | 1 | 1 |
| 3 div | 1 | 1 | 1 | 1 | 1 | 1 | 1 |
| 3 div+IRF | 1 | 1 | 1 | 1 | 1 | 1 | 1 |
| 4 div | 1 | 1 | 1 | 1 | 1 | 1 | 1 |
| 4 div+IRF | 1 | 1 | 1 | 1 | 1 | 1 | 1 |
| multiexponential 1.6 ns / 3.2 ns | | | | | | | |
| $1x10^1 s^{-1}$ | $1x10^1 s^{-1}$ | $1x10^2 s^{-1}$ | $2x10^2 s^{-1}$ | $5x10^2 s^{-1}$ | $1x10^3 s^{-1}$ | $5x10^3 s^{-1}$ | $1x10^4 s^{-1}$ |
| 1 div | 2 | 2 | 1 | 1 | 1 | 1 | 1 |
| 1 div+IRF | 2 | 1 | 1 | 1 | 1 | 1 | 1 |
| 2 div | 2 | 1 | 1 | 1 | 1 | 1 | 1 |
| 2 div+IRF | 2 | 1 | 1 | 1 | 1 | 1 | 1 |
| 3 div | 2 | 3 | 1 | 1 | 1 | 1 | 1 |
| 3 div+IRF | 2 | 1 | 3 | 1 | 1 | 1 | 1 |
| 4 div | 2 | 1 | 3 | 1 | 1 | 1 | 1 |
| 4 div+IRF | 3 | 1 | 1 | 1 | 1 | 1 | 1 |
| $1x10^2 s^{-1}$ | $1x10^1 s^{-1}$ | $1x10^2 s^{-1}$ | $2x10^2 s^{-1}$ | $5x10^2 s^{-1}$ | $1x10^3 s^{-1}$ | $5x10^3 s^{-1}$ | $1x10^4 s^{-1}$ |
| 1 div | 2 | 2 | 1 | 1 | 1 | 1 | 1 |
| 1 div+IRF | 1 | 2 | 2 | 1 | 1 | 1 | 1 |
| 2 div | 2 | 2 | 2 | 1 | 1 | 1 | 1 |
| 2 div+IRF | 2 | 2 | 2 | 1 | 1 | 1 | 1 |
| 3 div | 2 | 2 | 2 | 1 | 1 | 1 | 1 |
| 3 div+IRF | 1 | 2 | 2 | 1 | 1 | 1 | 1 |
| 4 div | 2 | 2 | 2 | 1 | 1 | 1 | 1 |
| 4 div+IRF | 2 | 2 | 3 | 1 | 1 | 1 | 1 |
| $2x10^2 s^{-1}$ | $1x10^1 s^{-1}$ | $1x10^2 s^{-1}$ | $2x10^2 s^{-1}$ | $5x10^2 s^{-1}$ | $1x10^3 s^{-1}$ | $5x10^3 s^{-1}$ | $1x10^4 s^{-1}$ |
| 1 div | 1 | 2 | 1 | 1 | 1 | 1 | 1 |
| 1 div+IRF | 1 | 2 | 1 | 1 | 1 | 1 | 1 |
| 2 div | 1 | 2 | 2 | 1 | 1 | 1 | 1 |
| 2 div+IRF | 2 | 2 | 2 | 1 | 1 | 1 | 1 |
| 3 div | 1 | 2 | 2 | 1 | 1 | 1 | 1 |
| 3 div+IRF | 1 | 2 | 2 | 1 | 1 | 1 | 1 |
| 4 div | 1 | 2 | 2 | 1 | 1 | 1 | 1 |
| 4 div+IRF | 1 | 2 | 2 | 1 | 1 | 1 | 1 |

| | | | | | | | |
|---|---|---|---|---|---|---|---|
| $5x10^2\ s^{-1}$ | $1x10^1\ s^{-1}$ | $1x10^2\ s^{-1}$ | $2x10^2\ s^{-1}$ | $5x10^2\ s^{-1}$ | $1x10^3\ s^{-1}$ | $5x10^3\ s^{-1}$ | $1x10^4\ s^{-1}$ |
| 1 div | 1 | 1 | 1 | 1 | 1 | 1 | 1 |
| 1 div+IRF | 1 | 1 | 1 | 1 | 1 | 1 | 1 |
| 2 div | 1 | 1 | 1 | 1 | 1 | 1 | 1 |
| 2 div+IRF | 1 | 1 | 1 | 1 | 1 | 1 | 1 |
| 3 div | 1 | 2 | 1 | 1 | 1 | 1 | 1 |
| 3 div+IRF | 1 | 2 | 1 | 1 | 1 | 1 | 1 |
| 4 div | 1 | 2 | 1 | 1 | 1 | 1 | 1 |
| 4 div+IRF | 1 | 2 | 1 | 1 | 1 | 1 | 1 |
| $1x10^3\ s^{-1}$ | $1x10^1\ s^{-1}$ | $1x10^2\ s^{-1}$ | $2x10^2\ s^{-1}$ | $5x10^2\ s^{-1}$ | $1x10^3\ s^{-1}$ | $5x10^3\ s^{-1}$ | $1x10^4\ s^{-1}$ |
| 1 div | 1 | 1 | 1 | 1 | 1 | 1 | 1 |
| 1 div+IRF | 1 | 1 | 1 | 1 | 1 | 1 | 1 |
| 2 div | 1 | 1 | 1 | 1 | 1 | 1 | 1 |
| 2 div+IRF | 1 | 1 | 1 | 1 | 1 | 1 | 1 |
| 3 div | 1 | 1 | 1 | 1 | 1 | 1 | 1 |
| 3 div+IRF | 1 | 1 | 1 | 1 | 1 | 1 | 1 |
| 4 div | 1 | 1 | 1 | 1 | 1 | 1 | 1 |
| 4 div+IRF | 1 | 1 | 1 | 1 | 1 | 1 | 1 |
| $5x10^3\ s^{-1}$ | $1x10^1\ s^{-1}$ | $1x10^2\ s^{-1}$ | $2x10^2\ s^{-1}$ | $5x10^2\ s^{-1}$ | $1x10^3\ s^{-1}$ | $5x10^3\ s^{-1}$ | $1x10^4\ s^{-1}$ |
| 1 div | 1 | 1 | 1 | 1 | 1 | 1 | 1 |
| 1 div+IRF | 1 | 1 | 1 | 1 | 1 | 1 | 1 |
| 2 div | 1 | 1 | 1 | 1 | 1 | 1 | 1 |
| 2 div+IRF | 1 | 1 | 1 | 1 | 1 | 1 | 1 |
| 3 div | 1 | 1 | 1 | 1 | 1 | 1 | 1 |
| 3 div+IRF | 1 | 1 | 1 | 1 | 1 | 1 | 1 |
| 4 div | 1 | 1 | 1 | 1 | 1 | 1 | 1 |
| 4 div+IRF | 1 | 1 | 1 | 1 | 1 | 1 | 1 |
| $1x10^4\ s^{-1}$ | $1x10^1\ s^{-1}$ | $1x10^2\ s^{-1}$ | $2x10^2\ s^{-1}$ | $5x10^2\ s^{-1}$ | $1x10^3\ s^{-1}$ | $5x10^3\ s^{-1}$ | $1x10^4\ s^{-1}$ |
| 1 diva | 1 | 1 | 1 | 1 | 1 | 1 | 1 |
| 1 div+IRF | 1 | 1 | 1 | 1 | 1 | 1 | 1 |
| 2 div | 1 | 1 | 1 | 1 | 1 | 1 | 1 |
| 2 div+IRF | 1 | 1 | 1 | 1 | 1 | 1 | 1 |
| 3 div | 1 | 1 | 1 | 1 | 1 | 1 | 1 |
| 3 div+IRF | 1 | 1 | 1 | 1 | 1 | 1 | 1 |
| 4 div | 1 | 1 | 1 | 1 | 1 | 1 | 1 |
| 4 div+IRF | 1 | 1 | 1 | 1 | 1 | 1 | 1 |
| multiexponential 1.6 ns / 3.6 ns | | | | | | | |

| 1x10$^{1}$ s$^{-1}$ | 1x10$^{1}$ s$^{-1}$ | 1x10$^{2}$ s$^{-1}$ | 2x10$^{2}$ s$^{-1}$ | 5x10$^{2}$ s$^{-1}$ | 1x10$^{3}$ s$^{-1}$ | 5x10$^{3}$ s$^{-1}$ | 1x10$^{4}$ s$^{-1}$ |
|---|---|---|---|---|---|---|---|
| 1 div | 2 | 2 | 1 | 1 | 1 | 1 | 1 |
| 1 div+IRF | 2 | 2 | 1 | 1 | 1 | 1 | 1 |
| 2 div | 2 | 3 | 1 | 1 | 1 | 1 | 1 |
| 2 div+IRF | 2 | 2 | 1 | 1 | 1 | 1 | 1 |
| 3 div | 2 | 2 | 1 | 1 | 1 | 1 | 1 |
| 3 div+IRF | 2 | 2 | 1 | 1 | 1 | 1 | 1 |
| 4 div | 2 | 2 | 1 | 1 | 1 | 1 | 1 |
| 4 div+IRF | 2 | 2 | 1 | 1 | 1 | 1 | 1 |

| 1x10$^{2}$ s$^{-1}$ | 1x10$^{1}$ s$^{-1}$ | 1x10$^{2}$ s$^{-1}$ | 2x10$^{2}$ s$^{-1}$ | 5x10$^{2}$ s$^{-1}$ | 1x10$^{3}$ s$^{-1}$ | 5x10$^{3}$ s$^{-1}$ | 1x10$^{4}$ s$^{-1}$ |
|---|---|---|---|---|---|---|---|
| 1 div | 2 | 2 | 2 | 1 | 1 | 1 | 1 |
| 1 div+IRF | 2 | 2 | 2 | 1 | 1 | 1 | 1 |
| 2 div | 2 | 2 | 2 | 1 | 1 | 1 | 1 |
| 2 div+IRF | 2 | 2 | 2 | 2 | 1 | 1 | 1 |
| 3 div | 2 | 2 | 2 | 2 | 1 | 1 | 1 |
| 3 div+IRF | 2 | 2 | 2 | 2 | 1 | 1 | 1 |
| 4 div | 2 | 2 | 2 | 2 | 1 | 1 | 1 |
| 4 div+IRF | 2 | 2 | 2 | 2 | 1 | 1 | 1 |

| 2x10$^{2}$ s$^{-1}$ | 1x10$^{1}$ s$^{-1}$ | 1x10$^{2}$ s$^{-1}$ | 2x10$^{2}$ s$^{-1}$ | 5x10$^{2}$ s$^{-1}$ | 1x10$^{3}$ s$^{-1}$ | 5x10$^{3}$ s$^{-1}$ | 1x10$^{4}$ s$^{-1}$ |
|---|---|---|---|---|---|---|---|
| 1 div | 1 | 2 | 2 | 1 | 1 | 1 | 1 |
| 1 div+IRF | 1 | 2 | 2 | 1 | 1 | 1 | 1 |
| 2 div | 1 | 2 | 2 | 2 | 1 | 1 | 1 |
| 2 div+IRF | 2 | 2 | 2 | 2 | 1 | 1 | 1 |
| 3 div | 1 | 2 | 2 | 2 | 1 | 1 | 1 |
| 3 div+IRF | 1 | 2 | 2 | 2 | 1 | 1 | 1 |
| 4 div | 1 | 2 | 2 | 2 | 1 | 1 | 1 |
| 4 div+IRF | 1 | 2 | 2 | 2 | 1 | 1 | 1 |

| 5x10$^{2}$ s$^{-1}$ | 1x10$^{1}$ s$^{-1}$ | 1x10$^{2}$ s$^{-1}$ | 2x10$^{2}$ s$^{-1}$ | 5x10$^{2}$ s$^{-1}$ | 1x10$^{3}$ s$^{-1}$ | 5x10$^{3}$ s$^{-1}$ | 1x10$^{4}$ s$^{-1}$ |
|---|---|---|---|---|---|---|---|
| 1 div | 1 | 2 | 1 | 1 | 1 | 1 | 1 |
| 1 div+IRF | 1 | 2 | 2 | 1 | 1 | 1 | 1 |
| 2 div | 1 | 2 | 2 | 1 | 1 | 1 | 1 |
| 2 div+IRF | 1 | 2 | 2 | 1 | 1 | 1 | 1 |
| 3 div | 1 | 2 | 2 | 1 | 1 | 1 | 1 |
| 3 div+IRF | 1 | 2 | 2 | 1 | 1 | 1 | 1 |
| 4 div | 1 | 2 | 2 | 1 | 1 | 1 | 1 |
| 4 div+IRF | 1 | 2 | 2 | 1 | 1 | 1 | 1 |

| 1x10$^{3}$ s$^{-1}$ | 1x10$^{1}$ s$^{-1}$ | 1x10$^{2}$ s$^{-1}$ | 2x10$^{2}$ s$^{-1}$ | 5x10$^{2}$ s$^{-1}$ | 1x10$^{3}$ s$^{-1}$ | 5x10$^{3}$ s$^{-1}$ | 1x10$^{4}$ s$^{-1}$ |
|---|---|---|---|---|---|---|---|
| 1 div | 1 | 1 | 1 | 1 | 1 | 1 | 1 |
| 1 div+IRF | 1 | 1 | 1 | 1 | 1 | 1 | 1 |

| | | | | | | | |
|---|---|---|---|---|---|---|---|
| 2 div | 1 | 1 | 1 | 1 | 1 | 1 | 1 |
| 2 div+IRF | 1 | 1 | 1 | 1 | 1 | 1 | 1 |
| 3 div | 1 | 1 | 1 | 1 | 1 | 1 | 1 |
| 3 div+IRF | 1 | 1 | 1 | 1 | 1 | 1 | 1 |
| 4 div | 1 | 1 | 1 | 1 | 1 | 1 | 1 |
| 4 div+IRF | 1 | 1 | 1 | 1 | 1 | 1 | 1 |
| $5x10^3$ $s^{-1}$ | $1x10^1$ $s^{-1}$ | $1x10^2$ $s^{-1}$ | $2x10^2$ $s^{-1}$ | $5x10^2$ $s^{-1}$ | $1x10^3$ $s^{-1}$ | $5x10^3$ $s^{-1}$ | $1x10^4$ $s^{-1}$ |
| 1 div | 1 | 1 | 1 | 1 | 1 | 1 | 1 |
| 1 div+IRF | 1 | 1 | 1 | 1 | 1 | 1 | 1 |
| 2 div | 1 | 1 | 1 | 1 | 1 | 1 | 1 |
| 2 div+IRF | 1 | 1 | 1 | 1 | 1 | 1 | 1 |
| 3 div | 1 | 1 | 1 | 1 | 1 | 1 | 1 |
| 3 div+IRF | 1 | 1 | 1 | 1 | 1 | 1 | 1 |
| 4 div | 1 | 1 | 1 | 1 | 1 | 1 | 1 |
| 4 div+IRF | 1 | 1 | 1 | 1 | 1 | 1 | 1 |
| $1x10^4$ $s^{-1}$ | $1x10^1$ $s^{-1}$ | $1x10^2$ $s^{-1}$ | $2x10^2$ $s^{-1}$ | $5x10^2$ $s^{-1}$ | $1x10^3$ $s^{-1}$ | $5x10^3$ $s^{-1}$ | $1x10^4$ $s^{-1}$ |
| 1 div | 1 | 1 | 1 | 1 | 1 | 1 | 1 |
| 1 div+IRF | 1 | 1 | 1 | 1 | 1 | 1 | 1 |
| 2 div | 1 | 1 | 1 | 1 | 1 | 1 | 1 |
| 2 div+IRF | 1 | 1 | 1 | 1 | 1 | 1 | 1 |
| 3 div | 1 | 1 | 1 | 1 | 1 | 1 | 1 |
| 3 div+IRF | 1 | 1 | 1 | 1 | 1 | 1 | 1 |
| 4 div | 1 | 1 | 1 | 1 | 1 | 1 | 1 |
| 4 div+IRF | 1 | 1 | 1 | 1 | 1 | 1 | 1 |

[a] Transition rates in this column indicate the transition rate from the short lifetime state, other columns indicate from the long lifetime state

[b] Optimizations using divisor scheme with single divisor

[c] Optimization using divisor scheme with a divisor for IRF and one additional divisor

[d] Optimizations using divisor scheme with two divisors

[e] Optimization using divisor scheme with a divisor for IRF and two additional divisors

[f] Optimizations using divisor scheme with three divisors

[g] Optimization using divisor scheme with a divisor for IRF and three additional divisors

[h] Optimizations using divisor scheme with four divisors

i Optimization using divisor scheme with a divisor for IRF and four additional divisors
